# Synchronized long-read genome, methylome, epigenome, and transcriptome for resolving a Mendelian condition

**DOI:** 10.1101/2023.09.26.559521

**Authors:** Mitchell R. Vollger, Jonas Korlach, Kiara C. Eldred, Elliott Swanson, Jason G. Underwood, Yong-Han H. Cheng, Jane Ranchalis, Yizi Mao, Elizabeth E. Blue, Ulrike Schwarze, Katherine M. Munson, Christopher T. Saunders, Aaron M. Wenger, Aimee Allworth, Sirisak Chanprasert, Brittney L. Duerden, Ian Glass, Martha Horike-Pyne, Michelle Kim, Kathleen A. Leppig, Ian J. McLaughlin, Jessica Ogawa, Elisabeth A. Rosenthal, Sam Sheppeard, Stephanie M. Sherman, Samuel Strohbehn, Amy L. Yuen, University of Washington Center for Mendelian Genomics (UW-CMG), Undiagnosed Diseases Network (UDN), Thomas A. Reh, Peter H. Byers, Michael J. Bamshad, Fuki M. Hisama, Gail P. Jarvik, Yasemin Sancak, Katrina M. Dipple, Andrew B. Stergachis

## Abstract

Resolving the molecular basis of a Mendelian condition (MC) remains challenging owing to the diverse mechanisms by which genetic variants cause disease. To address this, we developed a synchronized long-read genome, methylome, epigenome, and transcriptome sequencing approach, which enables accurate single-nucleotide, insertion-deletion, and structural variant calling and diploid *de novo* genome assembly, and permits the simultaneous elucidation of haplotype-resolved CpG methylation, chromatin accessibility, and full-length transcript information in a single long-read sequencing run. Application of this approach to an Undiagnosed Diseases Network (UDN) participant with a chromosome X;13 balanced translocation of uncertain significance revealed that this translocation disrupted the functioning of four separate genes (*NBEA*, *PDK3*, *MAB21L1*, and *RB1*) previously associated with single-gene MCs. Notably, the function of each gene was disrupted via a distinct mechanism that required integration of the four ‘omes’ to resolve. These included nonsense-mediated decay, fusion transcript formation, enhancer adoption, transcriptional readthrough silencing, and inappropriate X chromosome inactivation of autosomal genes. Overall, this highlights the utility of synchronized long-read multi-omic profiling for mechanistically resolving complex phenotypes.

## Introduction

The diagnosis of Mendelian conditions is challenged by the ability to accurately detect pathogenic genetic variation and the ability to determine whether an identified genetic variant has a functional consequence. Recent advances in accurate long-read sequencing have markedly improved our ability to detect genetic variants^1–4^, yet our ability to determine whether an identified genetic variant has a functional consequence remains quite limited. This challenge is particularly acute when evaluating non-coding genetic variants owing to the number of non-coding variants harbored in each genome and the diverse mechanisms by which non-coding variants can cause disease. Multi-omic approaches have shown promise in resolving this challenge by integrating genomic information with functional information from the same sample, such as RNA-seq and CpG methylation information^5–7^. However, the broader adoption of these multi-omic approaches for resolving Mendelian conditions has remained limited, in part due to experimental and analytical shortcomings with current multi-omic approaches. Specifically, current multi-omic approaches are grounded in short-read sequencing, which limits their ability to comprehensively characterize the functional output of individual haplotypes within a sample. In addition, current approaches combine data from independently prepared samples, which necessitates batch effect corrections between each ‘ome’.

Accurate long-read sequencing has the potential to resolve these limitations through haplotype-phased epigenetic and transcriptomic information. Specifically, accurate long-read sequencing permits the simultaneous detection of CpG methylation information on each sequenced read^8^, which provides a direct evaluation of the impact of genetic variants on CpG methylation patterns^9–11^. In addition, long-read sequencing can be leveraged to enable the simultaneous identification of chromatin architectures^12–15^, such as the occupancy of nucleosomes, transcription factors, and accessible chromatin patches along each sequenced read. For example, with single-molecule chromatin fiber sequencing (Fiber-seq) prior to DNA extraction permeabilized cells are treated with a non-specific N6-methyladenine methyltransferase (m6A-MTase), which stencils the architecture of each chromatin fiber onto its underlying DNA molecule in the form of m6A-modified bases, which can be directly read out during standard Single Molecule, Real-Time (SMRT) sequencing. Furthermore, recent advances in Fiber-seq enable the co-identification of both accurate chromatin features^16^ and CpG methylation along each sequenced read^17^, providing genomic sequence, CpG methylation, and chromatin epigenetic information in the same sequencing run. Finally, long-read full-length transcript sequencing is emerging as a powerful tool for resolving the transcript impact of genetic variants^18, 19^, and recent cDNA concatenation workflows for processing full-length transcript data (MAS-Seq)^20^ provide an order-of-magnitude higher throughput identification of full-length transcript data.

We sought to leverage recent advances in both Fiber-seq and MAS-Seq to create a synchronized long-read multi-ome, which provides an accurate, haplotype-phased, long-read genome, CpG methylome, chromatin epigenome, and transcriptome without the need for tiered sampling or redundant sequencing (**Fig. 1a**). We benchmarked the performance of this synchronized multi-ome assay against Genome-in-a-Bottle (GIAB) samples with known ground truths. We then applied it to solve the genetic and molecular basis of a participant within the UDN, demonstrating how each ‘ome’ contributes to revealing the molecular basis of this individual’s disease in a single sequencing run.

**Figure 1.**
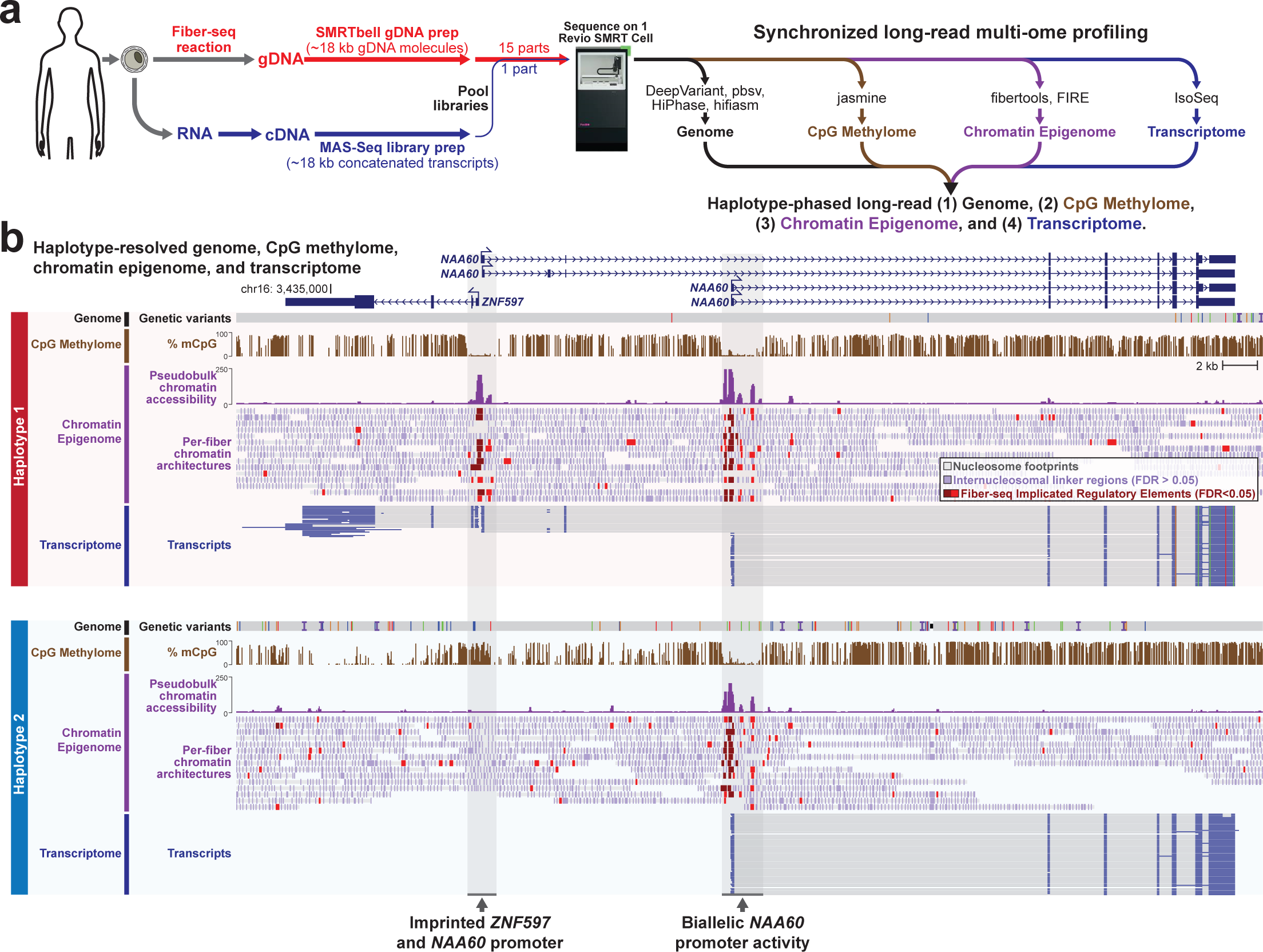
Synchronized long-read genome, methylome, epigenome and transcriptome sequencing. **a,** Schematic describing the experimental and computational workflow for synchronized multi-ome profiling. Specifically, cells are subjected to a Fiber-seq reaction followed by genomic DNA extraction and SMRTbell library preparation, and in parallel cells are subjected to an RNA extraction followed by complementary DNA (cDNA) synthesis and MAS-Seq library preparation. The two libraries are then mixed together and sequenced simultaneously using a single sequencing run, enabling the simultaneous detection of the genome, CpG methylome, chromatin epigenome, and transcriptome from the sample. **b,** Example genomic region showing the haplotype-resolved genome, CpG methylome, chromatin epigenome, and transcriptome from GM12878 cells at a known imprinted locus.

## Results

### Synchronized long-read multi-ome profiling

We first tested the accuracy of this approach for identifying genetic variants by applying long-read multi-ome profiling to two Genome-in-a-bottle (GIAB) cell lines that have been extensively characterized (GM12878 [HG001], and GM24385 [HG002]). Fiber-seq modified genomic DNA (gDNA) was generated by treating permeabilized cells with a non-specific m6A-MTase for 10 minutes, followed by gDNA extraction, shearing to ∼18 kb, and barcoding using standard SMRTbell library prep. In parallel, RNA was isolated from the same cells and polyA-primed cDNA was generated and concatenated into ∼18 kb molecules using a modified version of the MAS-Seq approach^20^ that we optimized for bulk RNA. Concatenating multiple cDNA molecules together into a ∼18 kb barcoded DNA molecule enables the gDNA and concatenated cDNA to have similar lengths, overcoming common issues that arise from pooling libraries of significantly different lengths into the same sequencing run (**Fig. 1a**). Pooling Fiber-seq gDNA and MAS-Seq concatenated cDNA at a molar ratio of 15:1 resulted in a total of over 6 million sequencing reads for each sample, with 5-10% of these reads deriving from MAS-Seq concatenated cDNA while the Fiber-seq gDNA reads permitted 26-30x genome-wide coverage (**Table 1**).

**Table 1.** Overview of genetic, epigenetic and transcript features of long-read multi-ome samples.

|  | Sample | HG001 | HG002 | UDN318336 |
| --- | --- | --- | --- | --- |
| **Sequencing Statistics:** | HiFi read yield | 98.8 Gb | 90.1 Gb | 90.3 Gb |
|  | HiFi reads | 6,268,744 | 6,000,613 | 5,902,290 |
|  | Insert length N50 | 18,932 bp | 17,443 bp | 17,221 bp |
|  | Median HiFi read QV | Q31 | Q32 | Q31 |
| **Genome:** | gDNA reads | 5,716,157 | 5,359,077 | 5,563,966 |
|  | gDNA read length N50 | 18,785 bp | 17,664 bp | 16,793 bp |
|  | Median coverage of hg38 | 30x | 26x | 28x |
|  | SNVs (F1) | 4,416,098 (99.90%) | 4,413,305 (99.94%) | 4,396,931 (n.a.) |
|  | Indels (F1) | 974,414 (98.75%) | 973,416 (98.81%) | 971,312 (n.a.) |
|  | SVs (F1) | 23,458 (n.a.) | 23,836 (92.5%) | 23,201 (n.a.) |
|  | % of reads haplotype phased | 87.0% | 90.0% | 87.7% |
|  | Assembly size (hap1/hap2) | 3,016 Mb / 3,018 Mb | 3,010 Mb / 3,010 Mb | 3,036 Mb / 3,038 Mb |
|  | Number of contigs (hap1/hap2) | 1,129 / 1,131 | 776 / 919 | 705 / 762 |
|  | Contig N50 (hap1/hap2) | 30.6 Mb / 30.7 Mb | 36.3 Mb / 28.8 Mb | 40.5 Mb / 27.3 Mb |
|  | QV (hap1/hap2) | 49.9 / 49.7 | 50.3 / 50.0 | n.a. |
| **mCpG Epigenome:** | Methylated CpG sites (%) | 12,818,712 (45%) | 20,252,720 (70%) | 20,301,742 (71%) |
|  | WGBS correlation (Pearson) | 0.79 | 0.92 | n.a. |
|  | ONT correlation (Pearson) | 0.82 | 0.93 | n.a. |
| **Chromatin Epigenome:** | Nucleosome size | 146 bp | 147 bp | 138 bp |
|  | Number of nucleosomes | 441,504,701 | 332,061,724 | 360,468,819 |
|  | FIREs in clusters* | 40% | 41% | 46% |
|  | FIREs in ENCODE peaks | 66% | 67% | 75% |
|  | Number of FIRE peaks† | 122,016 | 84,121 | 167,752 |
|  | % of FIRE peaks in ENCODE peaks | 91.6 | 97.3 | 96.3 |
| **Transcriptome:** | MAS-Seq reads | 518,921 | 604,272 | 297,248 |
|  | Transcript reads | 2,347,741 | 3,716,982 | 2,196,793 |
|  | Mean transcript length | 2,579 bp | 2,041 bp | 2,798 bp |
|  | Total unique genes | 10,536 | 11,512 | 10,383 |
|  | Total unique isoforms | 40,618 | 48,967 | 32,430 |

FIRE: Fiber-seq Implicated Regulatory Element
† Number of significant Fiber-seq peaks in the genome
\* Percent of FIREs preferentially clustered along the genome

### Genetic and epigenetic validation of synchronized long-read multi-ome profiling

Overall, this sequencing strategy did not detract from the ability to accurately identify genetic variants. Specifically, Fiber-seq modified gDNA from GM24385 cells retained highly accurate variant calling with a single nucleotide variant (SNV) F1 of 99.94%, an indel F1 of 98.81% and a structural variant (SV) F1 of 92.5% against established gold-standard variant annotations from HG002 (**Table 1**). In addition, as this approach is grounded in accurate long-read sequencing, 90.0% of Fiber-seq sequencing reads could be accurately phased to their respective haplotype, and these reads were compatible with generating high-quality long-read *de novo* diploid genome assemblies (haplotype contig N50s >20 Mb, consensus accuracy QV 50).

This approach also enabled the identification of epigenetic information. Specifically, CpG methylation information directly obtained using long-read multi-ome profiling correlated with existing whole genome bisulfite sequencing (WGBS) and Oxford Nanopore sequencing data (Pearson correlation of 0.92-0.93 for GM24385 cells) (**Table 1**). In addition, m6A modifications obtained using long-read multi-ome profiling demarcated ∼332 million nucleosome footprints (size ∼147 bp each). Using these single-molecule m6A-marked chromatin features, we identified 84,121 accessible elements within GM24385 cells, with 97.3% of these peaks mapping to known regulatory elements as defined by ENCODE^21^. Finally, this approach enabled the identification of over 2 million full-length non-concatenated transcripts, which arise from 48,967 unique splicing isoforms derived from 11,512 genes expressed in GM24385 cells. Overall, these data demonstrate strong agreement between synchronized long-read multi-ome profiling and that obtained using traditional short-read based transcript and epigenetic studies.

Long-read multi-ome profiling of these GIAB cells identified haplotype-phased genetic variation, CpG methylation, chromatin accessibility, and transcript data between both haplotype-phased genomes contained within these cells (**Fig. 1b and Extended Data Fig. 1**). For example, in GM12878 cells we identified 57 regulatory elements with significant haplotype-specific chromatin accessibility, many of which were contained within imprinted loci (**Extended Data Fig. 2**). Pairing the haplotype-resolved chromatin and transcriptomic data together at these loci enabled the identification of specific regulatory elements that were subjected to imprinting, as well as the transcriptional output of these regulatory elements (**Fig. 1b**). Furthermore, this approach identified non-coding genetic variants within challenging-to-map regions of the genome, such as the HLA locus, that appear to disrupt gene regulatory patterns (**Extended Data Fig. 3**).

### Long-read multi-ome profiling for identifying the molecular basis of a Mendelian condition

We next determined whether this approach could identify the molecular basis for a previously unsolved MC. Specifically, we applied synchronized long-read multi-ome profiling to skin fibroblasts from an undiagnosed 9-month-old female with bilateral retinoblastomas, developmental delay, polymicrogyria, sensorineural hearing loss (SNHL), lactic acidosis, hypotonia, and dysmorphic facial features who was enrolled within the UDN (UDN318336) (**Fig. 2a** and **Supplementary Table 1**). Prior trio genome sequencing did not reveal any genetic variants thought to be causing her phenotype, and a SNP array was normal (**Supplementary Table 2**). Clinical RNA sequencing of *RB1* revealed that bulk *RB1* expression was at the low end of normal, but *RB1* Sanger sequencing was normal. Karyotype revealed an apparently balanced *de novo* X;13 translocation (46,XX,t(X;13)(p22.1;q14.1)) of uncertain significance (**Fig. 2b**), and Fluorescence in situ hybridization (FISH) studies showed that *RB1* was unlikely to be disrupted at the breakpoint. Balanced translocations are common structural chromosomal rearrangements that result in copy neutral changes in the genome. While the majority of balanced translocations do not result in disease^22^, certain balanced translocations are known to cause Mendelian conditions when translocations disrupt genes, create fusion transcripts, alter gene regulatory programs, or involve the chromosome X, which can also result in the extension of X chromosome inactivation (XCI) into autosomal DNA^23^. These mechanisms often require an array of diverse functional studies to independently evaluate.

**Figure 2.**
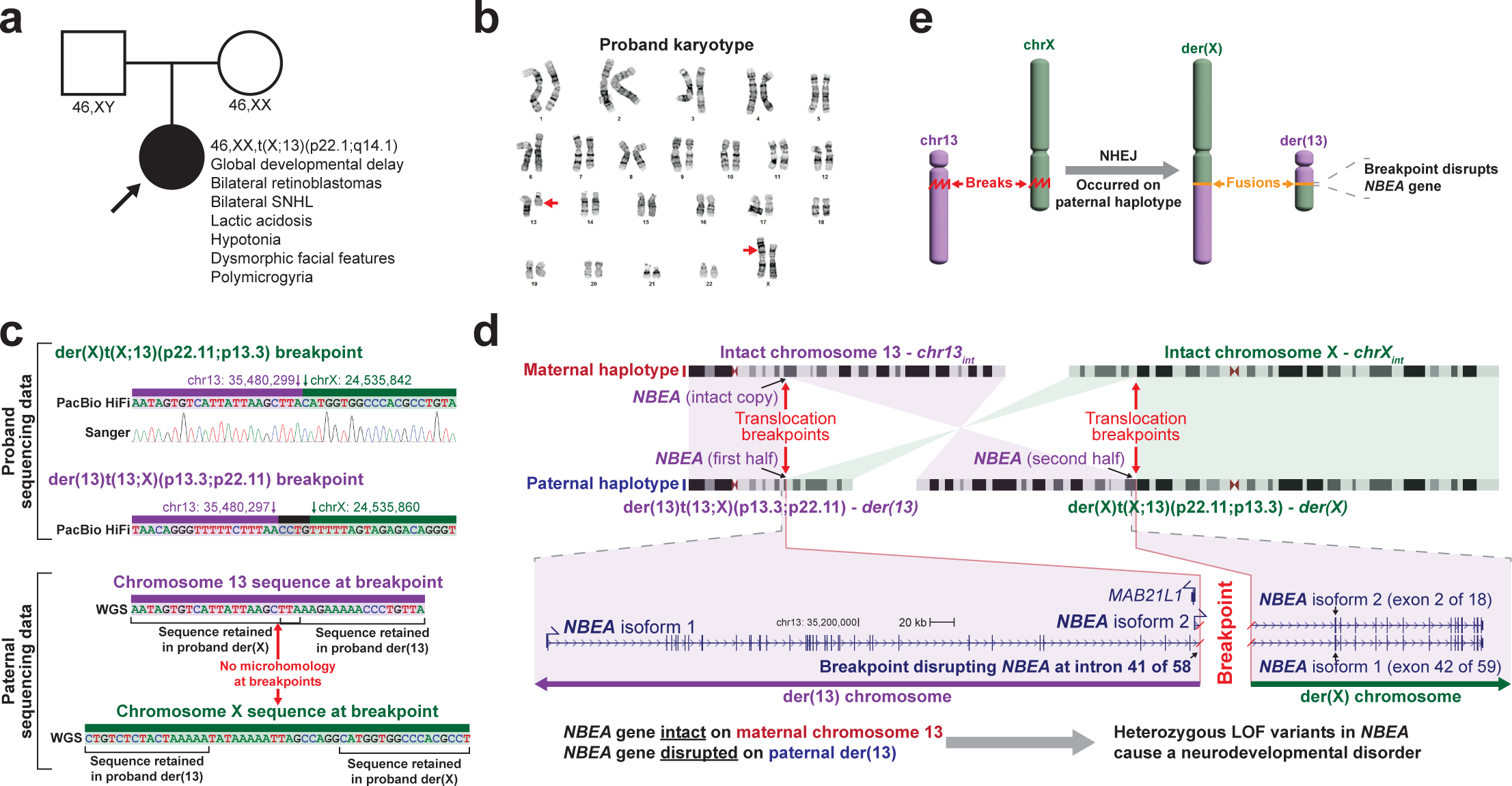
Long-read multi-ome for resolving the genetic basis of an unsolved Mendelian condition. **a,** Pedigree for the proband, as well as the clinical features of the proband and the results of her karyotype and that of her parents. **b,** Image of the proband’s karyotype with the der(13) and der(X) chromosomes marked by red arrows. **c,** Sequence of the breakpoints on der(X) and der(13), as well as the sequence of this same region in chromosomes 13 and X in her father. Sanger trace showing validation of the der(X) breakpoint junction. **d,** (top) Idiogram showing the intact chromosomes 13 and X, as well as the derivative chromosomes 13 and X in the proband. Translocation breakpoints, and the location of the gene *NBEA* are highlighted. (bottom) Gene model for both *NBEA* isoforms that differ in their transcriptional start site, showing the portion of *NBEA* that is located on der(13) versus der(X). **e,** Schematic showing the breakpoint and fusion event that occurred selectively on the paternal haplotype.

### Long-read genome exposes *NBEA* haploinsufficiency

Long-read multi-ome profiling of fibroblast cells from this individual resulted in 28x genome-wide haplotype-resolved coverage of the genome, CpG methylome, and chromatin epigenome, and additionally 2,196,793 full-length transcripts (**Table 1**). Genome assembly of these sequencing reads delineated the precise translocation breakpoints (**Fig. 2c**), which resulted in the creation of a ∼50 Mb derivative chromosome 13, der(13), that contains all of 13p, and part of 13q and Xp, as well as a ∼210 Mb derivative chromosome X, der(X), that contains all of Xq, and part of Xp and 13q. The translocation breakpoint on 13q is located in intron 41 of the 58-exon gene *NBEA*. Consequently, the first 41 exons of *NBEA* are located on der(13), and the terminal 17 exons are located on der(X) (**Fig. 2d**). The haplotype-phased multi-ome functional data demonstrated that although the *NBEA* promoters on the intact chromosome 13 (chr13_int_) and der(13) showed similar chromatin accessibility (**Extended Data Fig. 4**), full-length *NBEA* transcripts would only be able to arise from chr13_int_, consistent with this balanced translocation resulting in *NBEA* haploinsufficiency via nonsense-mediated decay of the transected *NBEA* transcripts (**Fig. 2e**).

### Long-read transcriptome exposes *PDK3-MAB21L1* fusion kinase transcript

*PDK3* on Xp encodes two isoforms that differ by the inclusion of a small terminal exon in isoform 2 that adds 7 amino acids to the protein sequence (**Fig. 3a**). The translocation breakpoint on Xp is located in this terminal intron for *PDK3* isoform 2, placing *PDK3* upstream of the gene *MAB21L1* with transcription for both of these genes in the same direction. The full-length transcript data revealed that this resulted in the formation of a fusion transcript between the splice donor site on exon 11 of *PDK3* isoform 2 and the splice acceptor site on exon 2 of *MAB21L1* isoform 2, adding 66 amino acids at the C-terminal end of PDK3. PDK3 is one of four kinases essential for phosphorylating and subsequently inhibiting pyruvate dehydrogenase complex (PDC) activity^24^. However, despite the production of this fusion kinase transcript, PDK3 protein levels remained low in these patient-derived fibroblast cells, consistent with low-level expression of *PDK3* in fibroblasts (**Extended Data Fig. 5**).

**Figure 3.**
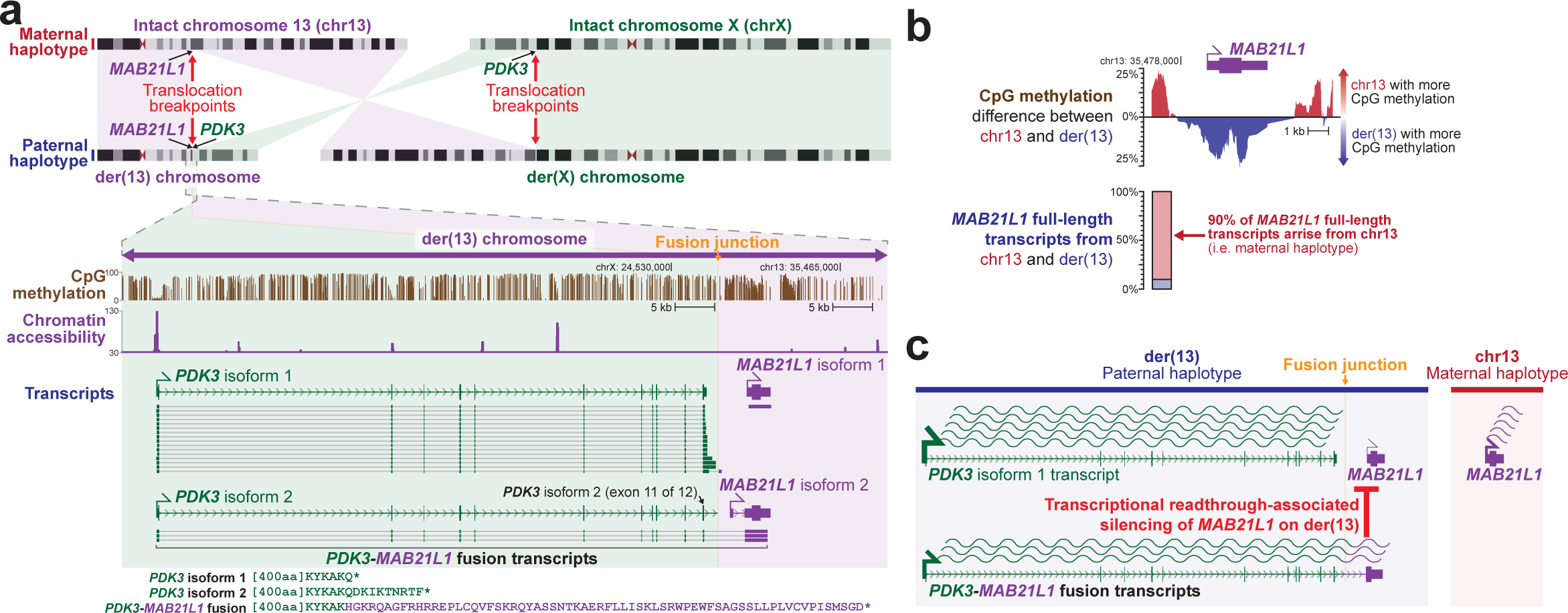
der(13) results in a *PDK3-MAB21L1* fusion transcript and *MAB21L1* silencing. **a,** (top) Idiogram showing the intact chromosomes 13 and X, as well as the derivative chromosomes 13 and X in the proband. Translocation breakpoints, and the location of the genes *PDK3* and *MAB21L1* are highlighted. (bottom) CpG methylation, chromatin accessibility, and full-length transcript data selectively on the der(13) haplotype are displayed, highlighting the formation of a fusion transcript between *PDK3* and *MAB21L1*. **b,** (top) CpG methylation differences at the *MAB21L1* promoter between chr13_int_ and der(13) demonstrating selective hyper-CpG methylation of the *MAB21L1* promoter along der(13). (below) Allelic imbalance of full-length *MAB21L1* transcripts between chr13_int_ and der(13) demonstrating silencing of *MAB21L1* along der(13). **c,** Schematic for transcriptional readthrough silencing of the *MAB21L1* gene selectively along der(13).

### Long-read CpG methylome exposes *MAB21L1* transcriptional readthrough silencing

Production of this fusion kinase transcript on der(13) results in transcriptional readthrough of the *MAB21L1* promoter. Transcriptional readthrough can result in complex epigenetic changes^25, 26^, including epigenetic silencing, as is seen at the *MSH2* promoter in individuals with *EPCAM* deletions^26^. Notably, we observed that the *MAB21L1* gene was epigenetically and transcriptionally silenced selectively along der(13), as demonstrated by allelic imbalance in both *MAB21L1* promoter CpG hypermethylation, and *MAB21L1* full-length transcripts (**Fig. 3b**). To validate this finding, we generated retinal organoids from these patient-derived fibroblasts, as *MAB21L1* is highly expressed in the retina (**Extended Data Fig. 6**). Fiber-seq on these patient-derived retinal organoids similarly demonstrated that the *MAB21L1* promoter is hypermethylated, and shows reduced chromatin accessibility selectively on der(13), consistent with transcriptional readthrough silencing of *MAB21L1* on der(13) (**Fig. 4a**).

**Figure 4.**
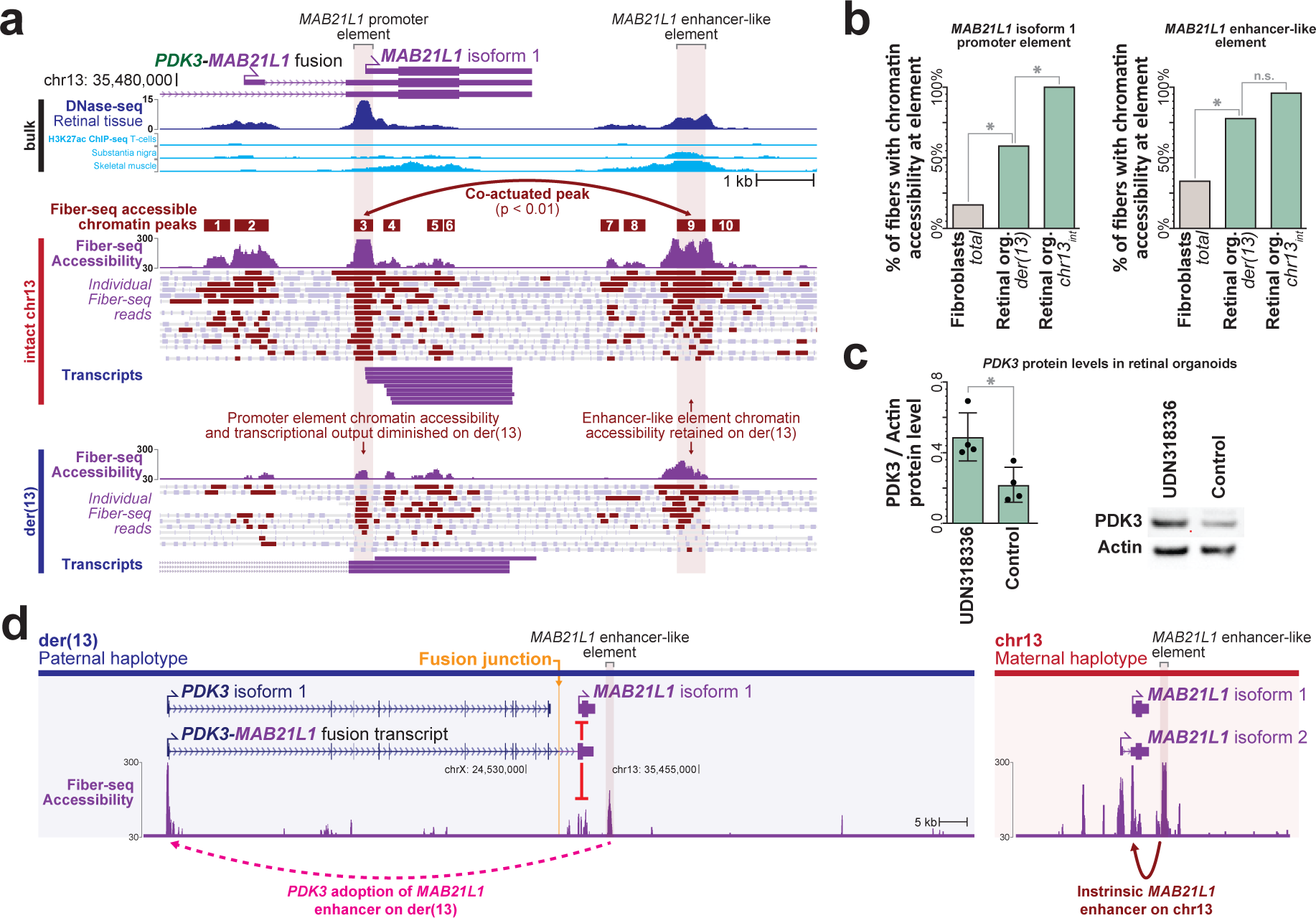
Placement of a *MAB21L1* enhancer-like element adjacent to *PDK3* along der(13) **a,** *MAB21L1* locus showing bulk DNase-seq and H3K27ac ChIP-seq (top), as well as haplotype-resolved Fiber-seq chromatin accessibility and full-length cDNA transcripts from patient-derived retinal organoids (bottom). Single chromatin accessibility peak pair with significant single-molecule co-actuated chromatin accessibility is shown, exposing a downstream enhancer-like element for *MAB21L1* isoform 1 promoter. **b,** Chromatin accessibility of the *MAB21L1* isoform 1 promoter, as well as the downstream enhancer-like element in patient-derived fibroblasts and retinal organoids (left), as well as their haplotype-specific accessibility in patient-derived retinal organoids (right). **c,** Bar plot and western blot showing PDK3 and beta-actin protein levels within patient-derived retinal organoids, as well as age-matched control retinal organoids. * p-value 0.0183 (T test) **d,** *MAB21L1* locus along the der(13) and chr13_int_ chromosomes showing the placement of a strong *MAB21L1* enhancer-like element in proximity to the *PDK3* promoter selectively along the der(13) haplotype.

### Long-read chromatin epigenome exposes *PDK3* enhancer adoption

Fiber-seq of the patient-derived retinal organoids also identified nine distinct accessible elements within 10 kb of the *MAB21L1* transcriptional start site, with element 9 demonstrating significant co-dependent actuation with the *MAB21L1* promoter, as well as bulk H3K27ac signal across multiple tissues (**Fig. 4a**), consistent with it being a *MAB21L1* enhancer. Notably, the accessibility of element 9 is selectively increased within the retinal organoids and element 9 remains accessible on 76% of the chromatin fibers deriving from der(13) in these cells, indicating that element 9 is a cell-selective *MAB21L1* enhancer that escapes transcriptional readthrough silencing (**Fig. 4b**), likely owing to its position after the transcription termination signal for the *PDK3-MAB21L1* fusion transcript. As the translocation positions this putative enhancer adjacent to the *PDK3* promoter, which is located only ∼70 kb away on der(13), we sought to see if the *PDK3* promoter is adopting this cell-selective *MAB21L1* enhancer. Consistent with this model of enhancer adoption, we find that patient-derived retinal organoids demonstrated significantly elevated PDK3 protein levels compared to control retinal organoids (**Fig. 4c** and **Extended Data Fig. 7**). Consequently, the chrX;13 translocation appears to simultaneously result in *NBEA* haploinsufficiency, a *PDK3-MAB21L1* fusion kinase transcript, transcriptional readthrough silencing of *MAB21L1*, and cell-selective enhanced *PDK3* expression via adoption of *MAB21L1* regulatory elements (**Fig. 4d**) - four distinct perturbations to three genes known to be involved in Mendelian conditions.

### Long-read multi-ome exposes inappropriate XCI of autosomal genes

*RB1* is the only gene associated with hereditary bilateral retinoblastomas, yet *RB1* is located 13.5 Mb away from the breakpoint along der(X). A chromosome X;13 translocation involving the opposite arm of chromosome X has previously been identified in an individual with retinoblastomas and was found to result in the inappropriate inactivation of the *RB1* locus in ∼10% of cells^23^ (**Extended Data Fig. 8**). As such, we sought to evaluate the haplotype-phased epigenetic data from this patient to determine whether der(X) is being subjected to XCI, and if this extended into autosomal DNA along der(X). Overall, we observed that patient-derived fibroblasts demonstrated markedly skewed XCI, with the intact chromosome X (chrX_int_) being preferentially subjected to XCI (**Fig. 5a**), consistent with prior observations in autosome-X translocations^27^. However, the autosomal DNA portion of der(X) did exhibit features of being epigenetically silenced in a subset of cells (**Fig. 5b**). Specifically, the 13p13.3 to 13qter region of chromosome 13, which is located on both der(X) and chr13_int_, selectively and significantly exhibited an imbalance in chromatin accessibility consistent with this region on der(X) being silenced in ∼3% of cells (**Fig. 5c**). Notably, this imbalance in chromatin accessibility was not seen in the two GIAB samples (**Extended Data Fig. 9a,b**), consistent with this being a feature driven by the balanced X;13 translocation. Consequently, in ∼3% of cells from this patient, the *RB1* locus would be epigenetically silenced (**Fig. 5d**).

**Figure 5.**
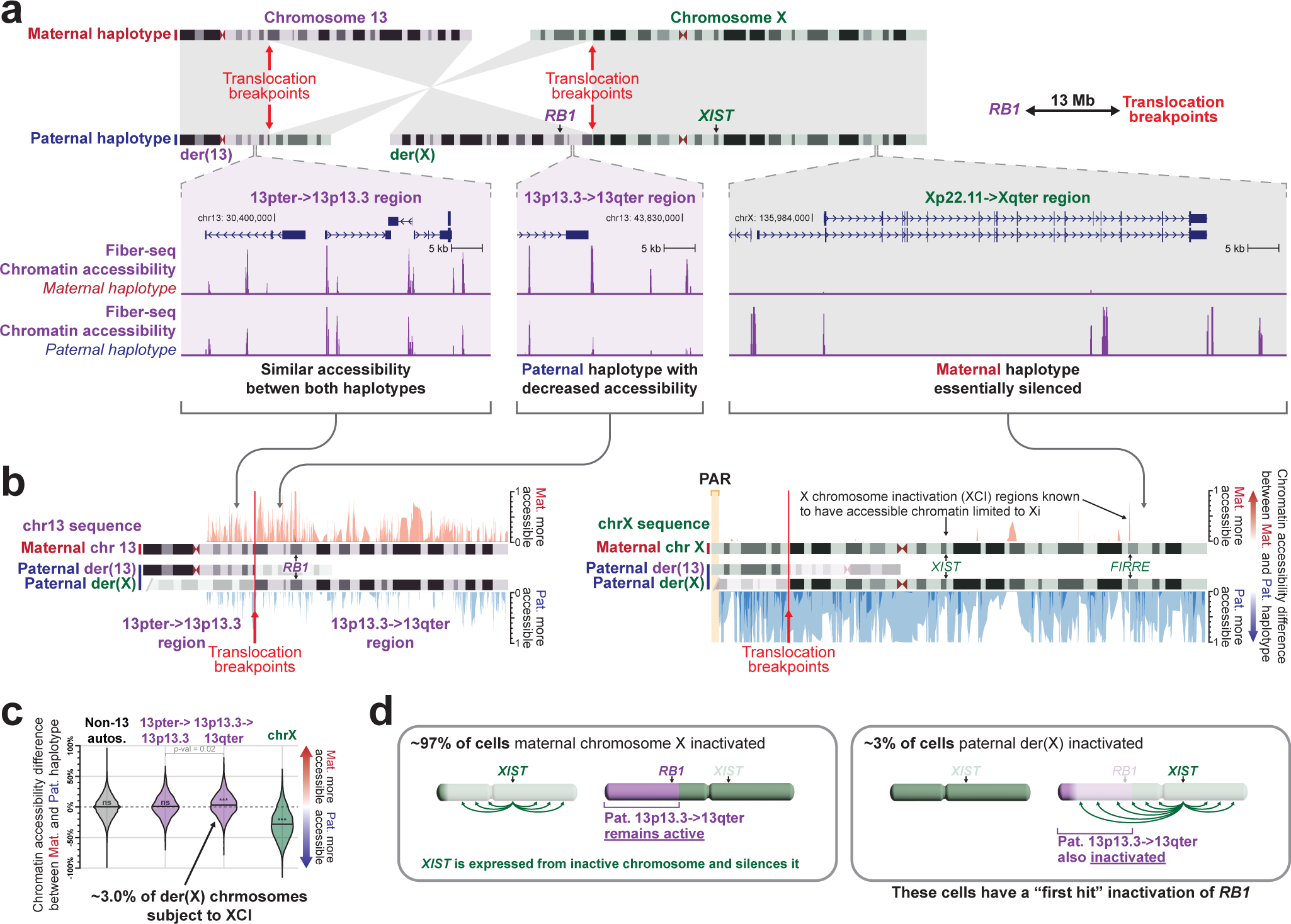
Inappropriate X chromosome inactivation of the *RB1* locus along der(X) **a,** (top) Idiogram showing the intact chromosomes 13 and X, as well as the derivative chromosomes 13 and X in the proband. Translocation breakpoints, and the location of the genes *RB1* and *XIST* are highlighted. (bottom) Haplotype-resolved chromatin accessibility is displayed for loci across the der(13) and der(X) chromosomes. **b,** Difference in chromatin accessibility between the maternal and paternal haplotype across loci along chromosome 13 (left) and chromosome X (right). Regions with more red than blue signal have more chromatin accessibility along chr13_int_ versus der(13) or chrX_int_ versus der(X). **c,** Swarm plot showing the overall haplotype imbalance in chromatin accessibility along autosomes (except for chromosome 13), chromosome X, and two portions of chromosome 13. Specifically, the 13pter->13p13.3 region is present along chr13_int_ and der(13), whereas the 13p13.3->13qter region is present along chr13_int_ and der(X). P-value calculated using Mann-Whitney U-test. **d,** Model showing inappropriate XCI of the autosomal region along der(X) that contains the *RB1* locus as the first hit for the development of bilateral retinoblastomas in this patient.

## Discussion

We present a synchronized long-read genomic, epigenomic, and transcriptomic sequencing approach, and demonstrate the accuracy and utility of this approach for characterizing the mechanistic basis by which genetic variants cause disease. In addition to eliminating redundant sequencing and batch effects that are inherent to current multi-ome methods, this synchronized long-read multi-ome approach enables the haplotype phasing of ∼90% of all genetic and epigenetic features, as well as full-length transcripts that contain heterozygous variants. As such, this approach enables the comparison of epigenetic and transcript signals between the two haplotypes within diploid individuals, providing an internally controlled comparison to identify heterozygous or biallelic variants that alter the epigenome or transcriptome. As most Mendelian conditions are caused by heterozygous or biallelic variants, this internally controlled approach is well suited for resolving the functional impact of disease-associated non-coding variants, in addition to phasing them relative to surrounding coding variants.

Application of this approach to an individual with a presumed MC and a chromosome X;13 balanced translocation revealed that this translocation disrupted the functioning of four separate genes associated with MCs, with each gene being disrupted via a distinct mechanism. Whereas the genomic data were able to identify one of these genes as being disrupted, the paired epigenetic and transcript data were essential for identifying the disruptions to the other three genes, highlighting the pivotal role paired functional data can play when evaluating an individual with a presumed MC. Furthermore, by pairing CpG methylation, chromatin, and transcript information together, this approach enabled the identification of quite distinct disease mechanisms, including nonsense mediated decay, formation of fusion transcripts, enhancer adoption, transcriptional read-through silencing, and inappropriate X chromosome inactivation of autosomal DNA.

Finally, this study defines a novel MC mediated by a single genetic variant, a 46,XX,t(X;13)(p22.11;q13.3) balanced translocation. This single genetic variant disrupts the functioning of four separate genes that are independently associated with MCs (MIMs 614041, 619157, 618479, and 300905), and her phenotype represents both an additive effect of these individual MCs, as well as novel phenotypes that previously have not been associated with these genes (**Table 2**). Specifically, it is likely that her developmental delay is at least partially caused by the *NBEA* loss-of-function (LOF) event, as *NBEA* haploinsufficiency is known to cause an autosomal dominant neurodevelopmental disorder^28^ (MIM 619157). Although her translocation results in the readthrough transcriptional silencing of *MAB21L1* on der(X), heterozygous *MAB21L1* LOF variants are not known to cause disease^29, 30^, consistent with her phenotype having little resemblance to that seen in individuals with biallelic *MAB21L1* LOF variants, which causes cerebellar, ocular, craniofacial, and genital syndrome (MIM 601280).

**Table 2.** Overview of molecular variants identified in UDN318336 via multi-ome long-read sequencing.

| Molecular event | 'ome(s) required for identification | Associated clinical phenotype | Proposed mechanism | Overlapping Mendelian condition |
| --- | --- | --- | --- | --- |
| *NBEA* haploinsufficiency | Genome | Developmental delay | *NBEA* nonsense-mediated decay on der(13) | MIM 619157 |
| *PDK3-MAB21L1* fusion kinase transcript | Transcriptome | Polymicrogyria, sensorineural hearing loss, developmental delay, lactic acidosis, and hypotonia. | Overexpression of *PDK3* in tissue that endogenously expresses *MAB21L1*. Potentially altered regulation of *PDK3-MAB21L1* fusion protein product. | MIM 300905; MIM 312170 |
| *PDK3* adoption of *MAB21L1* enhancer and subsequent *PDK3* ectopic gain-of-expression | Chromatin epigenome |  |  |  |
| X chromosome inactivation of *RB1* locus | Chromatin epigenome | Bilateral retinoblastomas | 'First hit' in development of biallelic *RB1* LOF | MIM 180200 |
| Transcriptional readthrough silencing of *MAB21L1* | CpG methylome; Chromatin epigenome; Transcriptome | No impact on patient phenotype as only one *MAB21L1* haplotype impacted, with other haplotype demonstrating intact gene regulation. | N/A | MIM 618479 |

This patient’s bilateral retinoblastomas appears to result from the inactivation of the *RB1* locus on der(X) in ∼3% of her cells (an extension of local X-chromosome inactivation), that results in the heterozygous LOF of *RB1* (MIM 614041). This would serve as the ‘first hit’ in the progression to biallelic *RB1* LOF. In combination with the prior case report of an individual with bilateral retinoblastomas and a balanced 46,X,t(X;13)(q28;q14.1) translocation^23^, these findings demonstrate that chromosome 13;X translocations involving either the p or q arm of chromosome X can result in a predisposition to bilateral retinoblastomas. Notably, although the inactivation of the autosomal region of der(X) extended into the *RB1* locus, which is located 13.5 Mb away from the translocation breakpoint, not all of the der(X) appeared to be equally impacted by the spreading of XCI along der(X) (**Extended Data Fig. 9c**). This indicates that the location of the breakpoints on both X and 13 may have a significant role on the extent of autosomal DNA that is subjected to XCI, and as such the phenotypic outcomes of the translocation.

Finally, her unique combination of alterations to both the transcript and expression patterns of *PDK3* appear to define a novel condition that overlaps with both *PDK3* gain-of-function variants, as well as PDC loss-of-function variants. A gain-of-function variant in *PDK3* has previously been shown to cause X-linked dominant Charcot-Marie-Tooth disease (CMTX6)^31^ (MIM 300905), which is characterized by childhood onset muscle weakness, muscle atrophy, sensory abnormalities, and SNHL. This gain-of-function variant increases PDK3 affinity for the pyruvate dehydrogenase complex (PDC), locking PDC in an inactive state. In contrast, complete PDC deficiency (MIM 312170) is associated with lactic acidosis, developmental delay, hypotonia, seizures, polymicrogyria and dysmorphic features^32^, a phenotype that is more severe than that of CMTX6 likely owing to the tissue specific expression of PDK3 protein, and the presence in many tissues of other kinases that regulate PDC. Notably, this patient has aspects of both CMTX6 and PDC deficiency, including polymicrogyria, SNHL, developmental delay, lactic acidosis, and hypotonia. This is likely due to the unique molecular mechanisms driving PDK3 dysfunction in her. Specifically, she has a fusion PDK3 kinase protein, ectopic gain-of-expression (GOE) of *PDK3* via enhancer adoption of a *MAB21L1* enhancer, and skewed XCI resulting in the selective expression of this enhanced fusion *PDK3* kinase transcript. As the intrinsic expression of *MAB21L1* is largely limited to neural tissues, it is likely that this enhancer adoption is selectively upregulating *PDK3* in those tissues. Consequently, in addition to the cell types that intrinsically express *PDK3*, she would exhibit PDK3 dysfunction in the brain, which typically expresses *PDK3* at low levels, likely extending PDC inhibition to this tissue and causing a phenotypic expansion beyond that seen in CMTX6.

Overall, we demonstrate that synchronized long-read multi-ome sequencing enables both the highly accurate discovery of haplotype-phased genetic variants, in addition to the simultaneous identification of the functional consequence of these genetic variants. It is anticipated that this approach will enable clinicians and researchers to better understand how diverse classes of genetic variation mechanistically drive human diseases, as well as potential molecular targets for modulating these diseases.

---

**Extended Data Figure 1.**
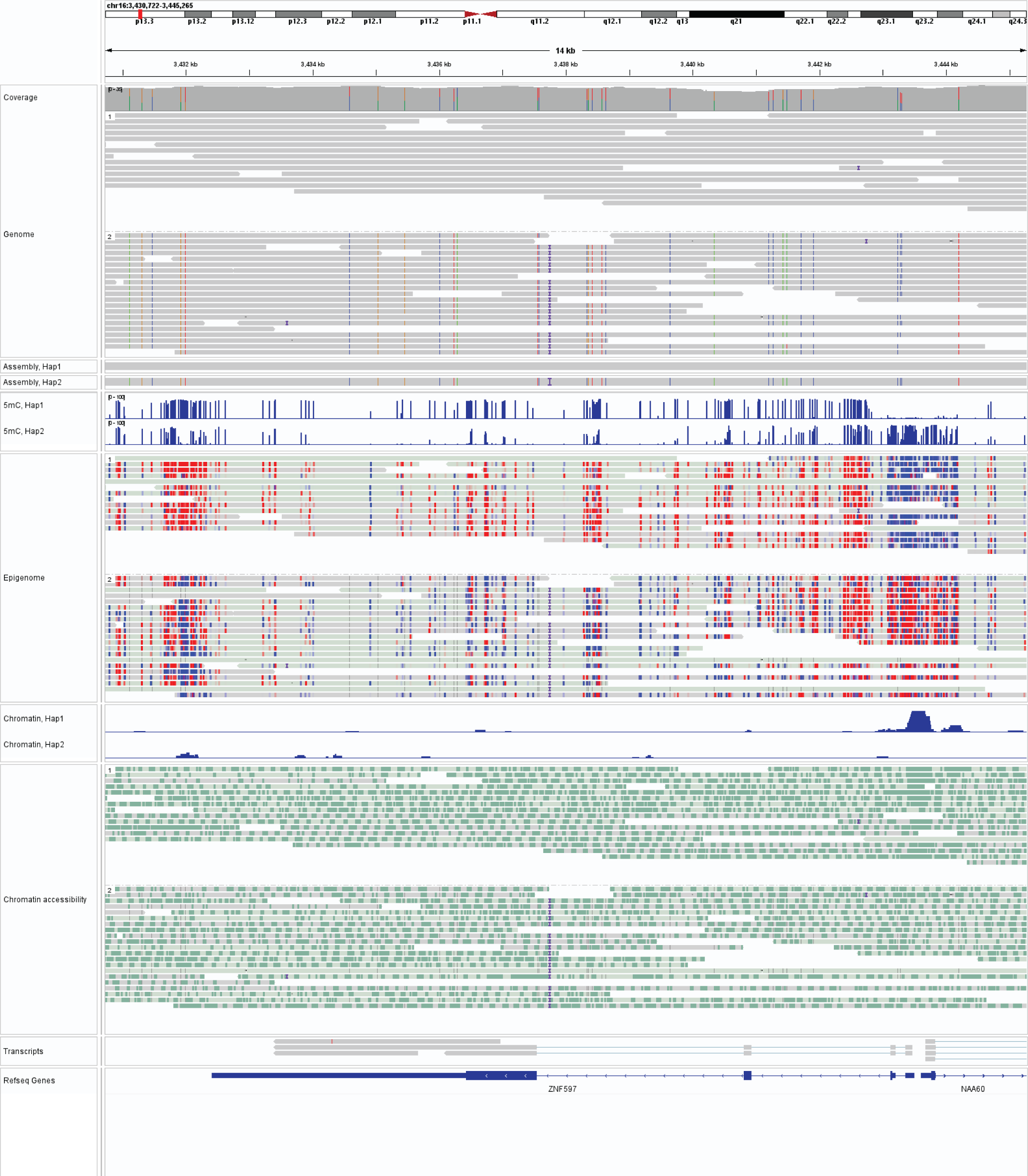
IGV view of integrated long-read multi-ome data. IGV view showing an example genomic region for GM12878 (from top to bottom panels) the haplotype-resolved genome, CpG methylome, chromatin, and full-length transcript annotations.

**Extended Data Figure 2.**
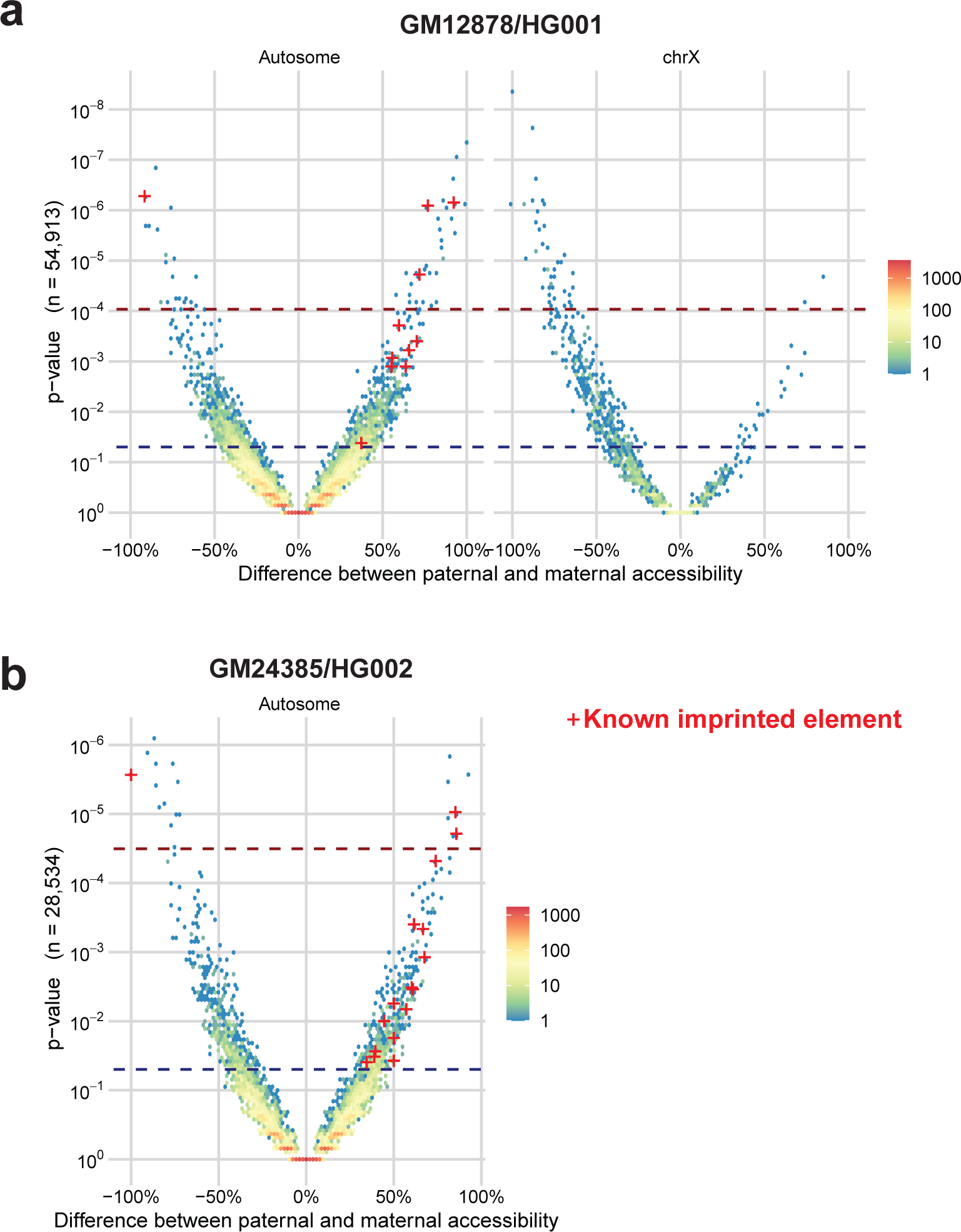
Haplotype-specific chromatin architectures. **a,** Volcano plot showing the absolute difference in the % of chromatin fibers with chromatin accessibility for each peak genome-wide between the paternal and maternal haplotypes for GM12878. Peaks are divided into whether they are present on an autosome, or the X chromosome. P-value was calculated using the Fisher exact test. Blue dash represents nominal significance line (p<0.01), red dash represents the Benjamini Hochberg FDR correction significance line. Peaks corresponding to known imprinted loci with nominally significant scores are denoted by red crosses. **b,** Same as a, but for GM24385. Data from the X chromosome is not shown, as GM24385 only has one haplotype for the X chromosome.

**Extended Data Figure 3.**
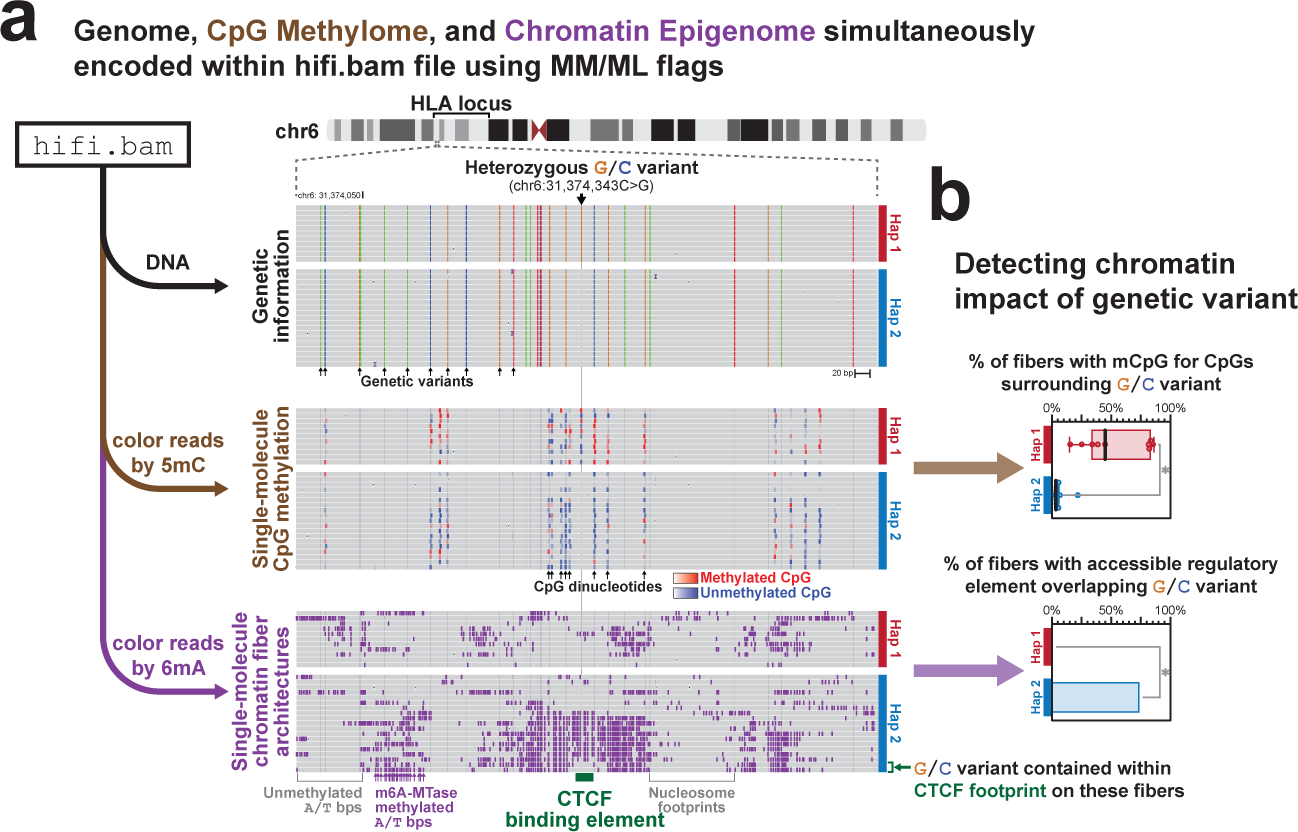
Identification of haplotype-specific chromatin architectures within HLA locus. **a,** View of single-molecule haplotype-resolved genetic information, CpG methylation information, and chromatin accessibility information for a heterozygous single-nucleotide variant (SNP) identified within the HLA locus. Note that all information is derived from the same sequencing reads. Denoted below is the location of a predicted CTCF binding element, and immediately above it are two fibers that demonstrate single-molecule protein occupancy at this site. **b,** Quantification of CpG methylation surrounding this SNP (top), as well as the % of fibers with a FIRE element overlapping that SNP (bottom), by haplotype. * p<0.01 (Fisher exact test).

**Extended Data Figure 4.**
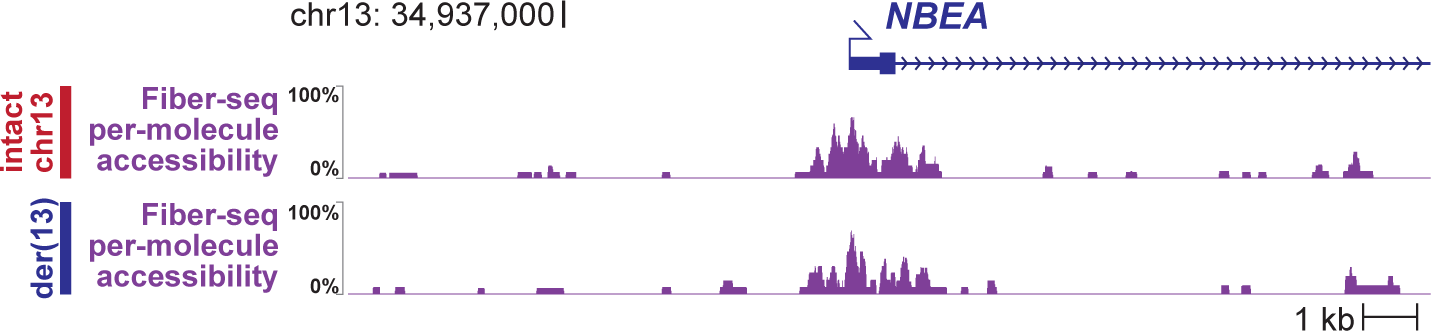
Haplotype-specific chromatin accessibility of *NBEA* promoter. Per-molecule chromatin accessibility of the NBEA promoter along both the der(13) and intact chr13 chromosomes within patient derived fibroblasts. Chromatin accessibility measured using Fiber-seq.

**Extended Data Figure 5.**
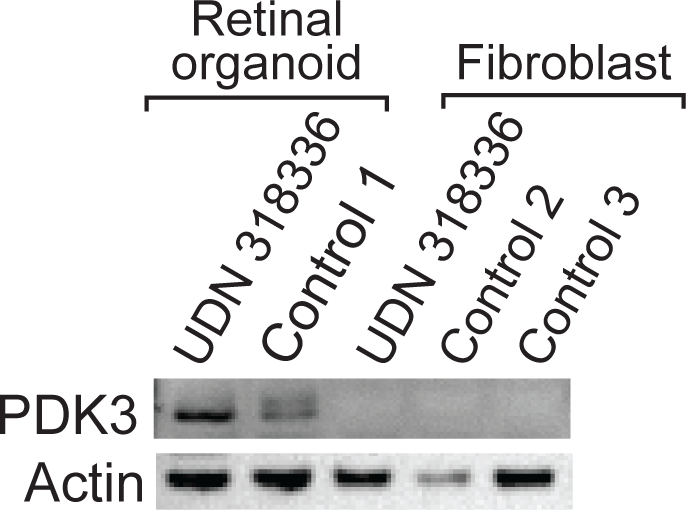
PDK3 protein levels in patient cells. Western blot of PDK3 and actin protein levels in patient derived fibroblasts and retinal organoids, as well as control age matched retinal organoids and fibroblast cultures.

**Extended Data Figure 6.**
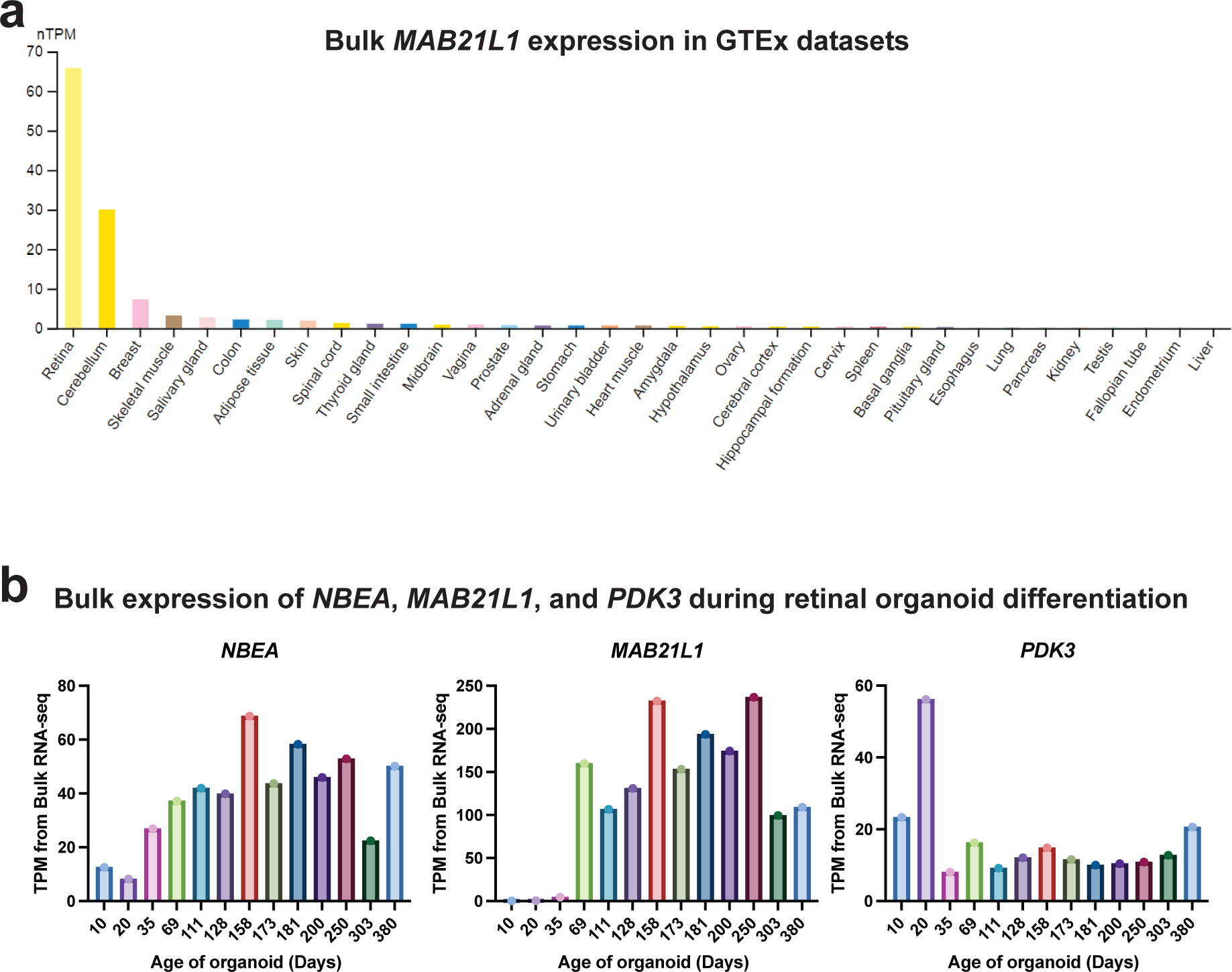
Cell-selective expression of *MAB21L1*. **a,** Barplot showing the bulk tissue RNA expression of *MAB21L1* (ENSG00000180660) from various GTEx samples are displayed by www.proteinatlas.org. **b,** Bulk expression of *NBEA*, *MAB21L1*, and *PDK3* during retinal organoid differentiation as a function of the age of the organoid. TPM - transcripts per million.

**Extended Data Figure 7.**
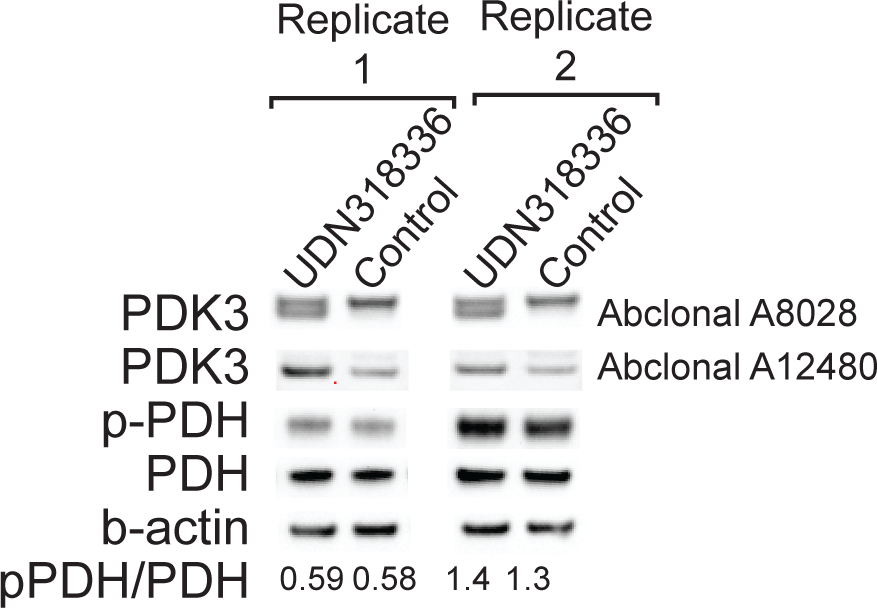
PDK3 protein levels in patient retinal organoids. Western blot images showing quantification of PDK3, phospho-PDH, PDH, and beta-actin levels in patient-derived retinal organoids, as well as age matched retinal organoids from a separate unaffected and unrelated individual. Data from two separate protein extractions are displayed. Quantification of phospho-PDH to PDH level is shown at bottom.

**Extended Data Figure 8.**
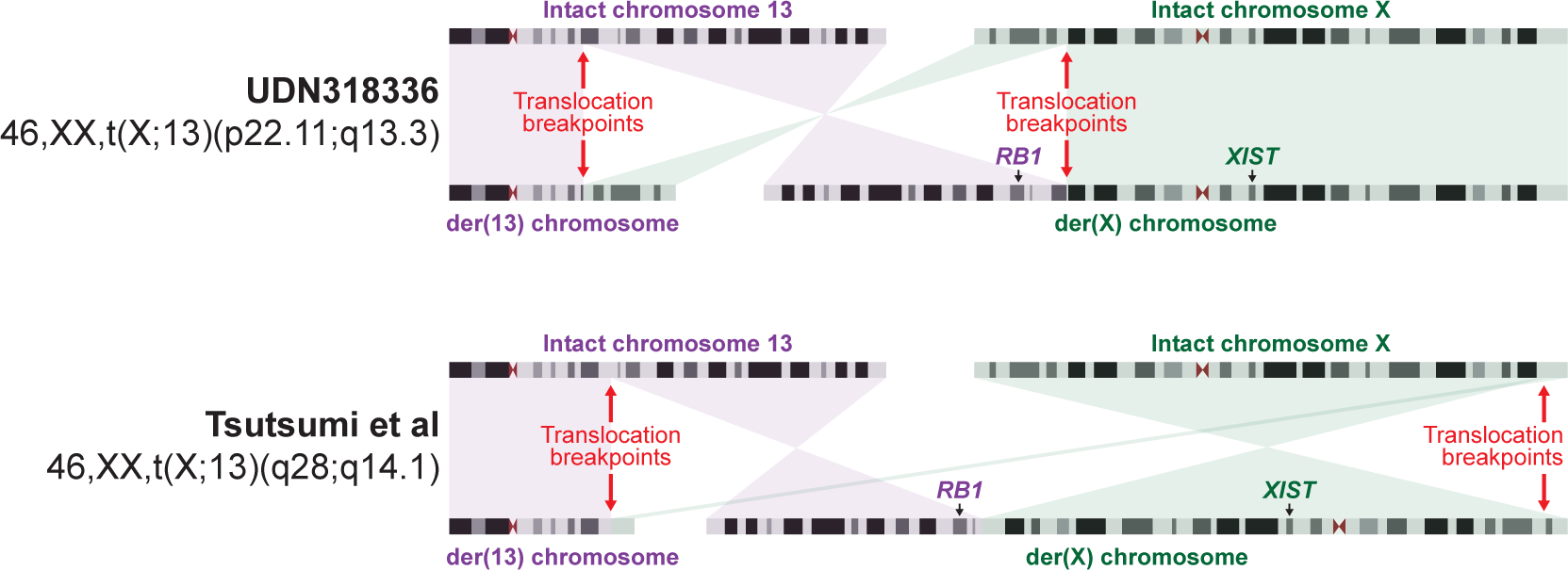
Comparison with prior chromosome X;13 translocations. Idiogram showing the translocation breakpoints and derivative chromosomes for this case, as well as a previously published case who similarly had bilateral retinoblastomas. The translocation breakpoints for the previously published case are estimated based on the karyotype.

**Extended Data Figure 9.**
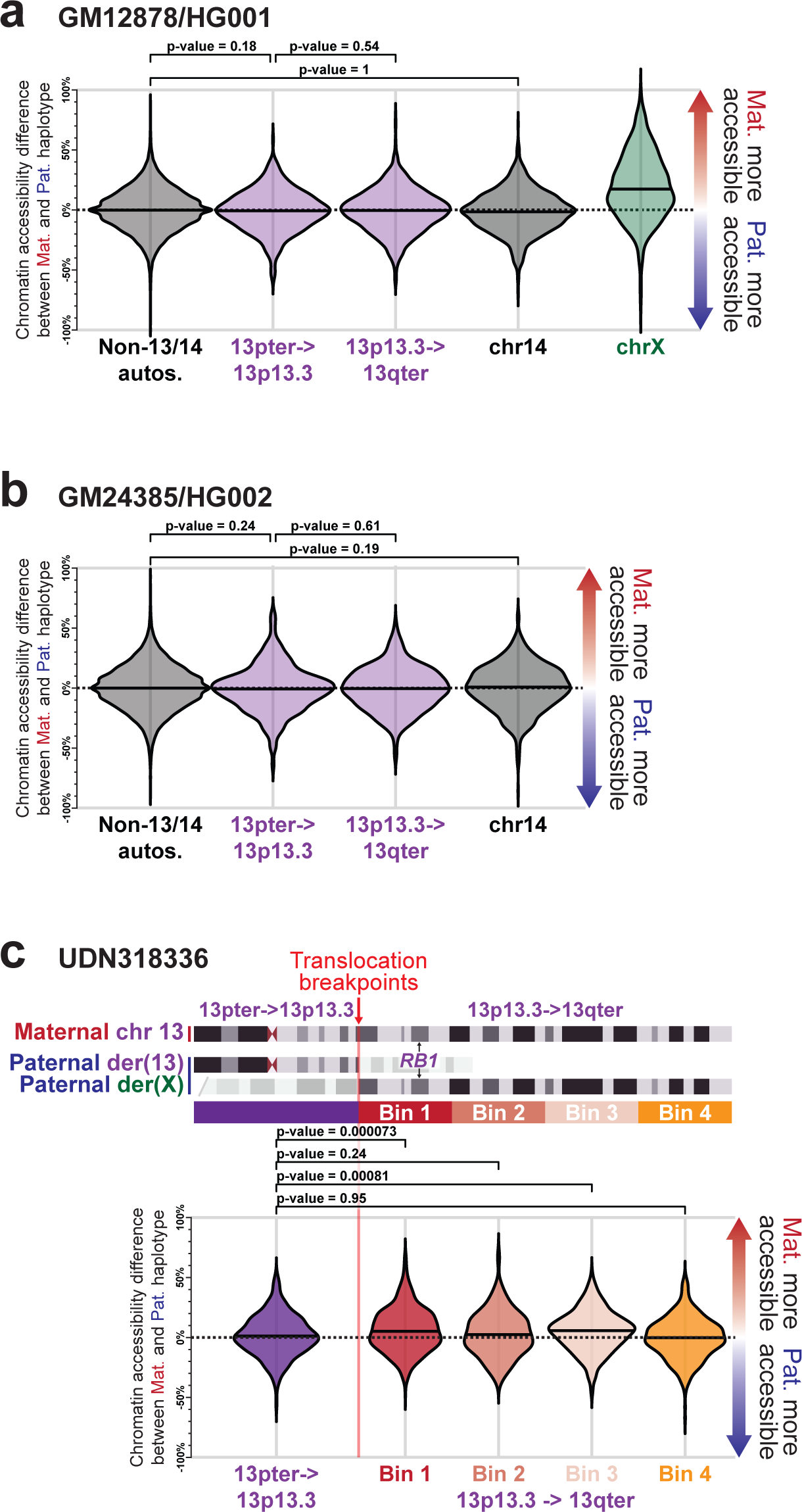
Allelic imbalance differences along chr13. **a,** Swarm plot showing the overall haplotype imbalance in chromatin accessibility along autosomes (except for chromosomes 13 and 14), chromosome X, chromosome 14, and two portions of chromosome 13 within GM12878. P-value calculated using Fisher exact test. **b,** Same as a, but for GM24385 cells. **c,** Swarm plot showing the haplotype imbalance in chromatin accessibility at different regions of chromosome 13 in fibroblasts from UDN318336.

## Methods

### Genomic DNA preparation

GM12878 (HG001) and GM24385 (HG002) cells were obtained as cells from Coriell. UDN318336 cultured fibroblast cells were obtained via a skin punch biopsy from participant UDN318336. UDN318336 retinal organoids were generated by first converting UDN318336 cultured fibroblast cells into induced pluripotent stem cells (iPSCs) using the CytoTune™-iPS Sendai reprogramming method (Invitrogen), and then differentiating these iPSCs into retinal organoids as previously described^33^. Retinal organoids were harvested at day 145 and then made into a single-cell suspension using Papain and DNAse for 20 min, then stopped with Ovomucoid (Worthington Biochemical LK003150) prior to permeabilization. Cells were permeabilized and treated with Hia5 enzyme as previously described^34^. Specifically, 1 million cells were washed with PBS and then resuspended in 60 μl Buffer A (15 mM Tris, pH 8.0; 15 mM NaCl; 60 mM KCl; 1mM EDTA, pH 8.0; 0.5 mM EGTA, pH 8.0; 0.5 mM Spermidine) and 60 μl of cold 2X Lysis buffer (0.1% IGEPAL CA-630 in Buffer A for GM12878 and GM24385; 0.2% IGEPAL CA-630 in Buffer A for UDN318336 fibroblasts and retinal organoid cells) was added and mixed by gentle flicking then kept on ice for 10 minutes. Samples were then pelleted, supernatant removed, and then resuspended in 57.5 μl Buffer A and moved to a 25°C thermocycler. 0.5 μl of Hia5 MTase (100 U) and 1.5 μl 32 mM S-adenosylmethionine (NEB B9003S) (0.8 mM final concentration) were added, then carefully mixed by pipetting the volume up and down 10 times with wide bore tips. The reactions were incubated for 10 minutes at 25°C then stopped with 3 μl of 20% SDS (1% final concentration) and transferred to new 1.5 mL microfuge tubes. High molecular weight DNA was then extracted using the Promega Wizard HMW DNA Extraction Kit A2920. PacBio SMRTbell libraries were then constructed using these Fiber-seq treated gDNA following the manufacturer’s SMRTbell prep kit 3.0 procedure (https://www.pacb.com/wp-content/uploads/Procedure-checklist-Preparing-whole-genome-and-metagenome-libraries-using-SMRTbell-prep-kit-3.0.pdf).

### RNA preparation

Bulk mRNA was isolated from GM24385 (HG002) cell cultures using the TRIzol Reagent and Phasemaker Tubes Complete System (Thermo Fisher). Bulk mRNA from GM12878 (HG001) and UDN318336 fibroblast RNA was isolated using the Qiagen RNeasy Plus Micro Kit (Catalog # 74034). RNA was subjected to cDNA synthesis as described in the standard Iso-Seq protocol (https://www.pacb.com/wp-content/uploads/Procedure-checklist-Preparing-Iso-Seq-libraries-using-SMRTbell-prep-kit-3.0.pdf), with the modification of using new cDNA amplification primers that combine the protocol’s sequence with MAS-Seq capture primer sequence^20^ to enable subsequent MAS-Seq concatenation. Following cDNA amplification, MAS-Seq libraries were generated by adapting the MAS-Seq protocol (https://www.pacb.com/wp-content/uploads/Procedure-checklist-preparing-MAS-Seq-libraries-using-MAS-Seq-for-10x-single-cell-3-kit.pdf) to an 8-mer array.

### Pooling Fiber-seq and MAS-Seq

Fiber-seq libraries and bulk MAS-Seq libraries from the same sample (i.e. GM12878, GM24385, or UDN318336 fibroblasts) were pooled in a 15:1 molar ratio, respectively, and sequenced using a single SMRT Cell on the PacBio Revio system. UDN318336 retinal organoid Fiber-seq was sequenced on a Sequel II system.

### Separating MAS-Seq and Fiber-seq reads

HiFi read bam files were computed and segregated into gDNA and MAS-Seq reads on the Revio system, taking advantage of the different adapters used in the two library protocols. MAS-Seq reads were de-concatenated into Iso-Seq reads using SMRT Link v12.0.

### Identification of genomic variants

Variant calling was performed with DeepVariant for SNVs and Indels^35^ (1.5.0) and pbsv for SVs (https://github.com/PacificBiosciences/pbsv). Reads were haplotype-phased using a custom pipeline (https://github.com/mrvollger/k-mer-variant-phasing) that uses SNVs identified with DeepVariant, and then runs a variant based phaser (HiPhase) to bin reads into phase blocks, which are assigned to either the maternal or paternal haplotype using parental short-read genome sequencing combined with the trio k-mer based phaser meryl. F1 statistics were calculated using GIAB variant calls from HG001 and HG002 as the gold standard. Specifically, for SNV and indel benchmarking we used the v4.2.1 GRCh38 benchmark for both NA12878_HG001 and HG002_NA24385_son. For SV benchmarking in HG002, we used the NIST_SV_v0.6 benchmark lifted to GRCh38 with NCBI Remap. *De novo* assembly was performed using hifiasm^36^, the concordance QVs of the assemblies against HG001 and HG002 GIAB references were computed with yak (https://github.com/lh3/yak).

### Identification of CpG methylation

Base-level CpG methylation was called using jasmine (PacBio). The percent CpG methylation at each genomic position was identified from a pileup of reads using pb-CpG-tools (https://github.com/PacificBiosciences/pb-CpG-tools). Pearson correlations for HG001 and HG002 were calculated using methylKit^37^, compared to whole genome bisulfite (WGBS) data from EpiQC^38^, and ONT data from Epi2me (https://labs.epi2me.io/giab-2023.05/, using the ‘super accuracy’ analysis level 60x bam files).

### Identification of chromatin architectures

m6A methylation and methyltransferase sensitive patches (MSPs) were identified using fibertools-rs^16^. MSPs correspond to regions between nucleosome footprints and represent a combination of internucleosomal linker regions, as well as accessible regulatory elements. MSPs were then processed using the Fiber-seq Inferred Regulatory Element (FIRE) pipeline (https://github.com/fiberseq/fiberseq-fire) (Vollger et al., in preparation). Briefly, the FIRE pipeline using a semi-supervised machine learning approach^39^ trained to identify the difference in signal between a regulatory element and an internucleosomal linker region. This method assigns an estimated precision based on validation data to every MSP, indicating the precision of the model for assigning a given MSP as a regulatory element versus an internucleosomal linker region. We then define significant MSPs (precision over 90%) to be Fiber-seq inferred regulatory elements (FIREs), and aggregate the FIRE signal over multiple Fiber-seq molecules to create a track representing genome-wide accessibility using the following formula:

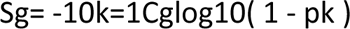

Where S is the aggregate FIRE signal at a genomic position (g), C is the genomic coverage of Fiber-seq data at that position, and p is the precision assigned to an individual FIRE. We then apply a genome-wide bonferroni correction (at alpha = 0.01) to call peaks within this signal track. Known imprinted loci were identified from^40^.

### Identification of full-length transcripts

Transcriptome statistics were generated using the Iso-Seq analysis workflow in SMRT Link v12.0. Fusion transcripts were identified using pbfusion (https://github.com/PacificBiosciences/pbfusion).

### Sanger validation of der(X) fusion junction

The der(X) fusion junction was amplified by PCR with primers annealing to sequences on chromosome 13 (5’-AAGCGTTCGAAATAACAGTGGC, chr13:35,480,163-35,480,184) and chromosome X (5’-ACCTCTGTGTTCCCAAAACCC, chrX:24,535,577-24,535,597)[hg38] that yielded a 403 bp amplicon. The product was checked by PAGE and sequenced in both directions with BigDye^TM^ Terminator v3.1 on the ABI 3500 Genetic Analyzer. The data were analyzed with Chromas software.

### *MAB21L1* expression

*MAB21L1* GTEx expression data was accessed from https://www.gtexportal.org/home/gene/MAB21L1. *NBEA*, *MAB21L1*, and *PDK3* expression during retinal organoid differentiation was quantified using publicly available data^41^. *MAB21L1* full-length transcripts from UDN318336 were haplotype-resolved using heterozygous variants contained within the transcript.

### Identification of *MAB21L1* putative enhancer peaks

Co-dependency scores for every neighboring FIRE peak with the *MAB21L1* promoter using UDN318336 retinal organoid Fiber-seq data was calculated as below. Specifically, we defined co-dependency scores as the observed rate of co-accessibility (i.e., accessible at both peaks along an individual chromatin fiber) minus the expected rate of co-accessibility given independence between the two peaks. Specifically, we identified overlapping chromatin fibers and accessible patches (FIRE precision ≤ 0.05) using bedtools intersect (v2.30.0) and calculated the proportion of fibers that are accessible at each peak. We performed all subsequent co-dependency analyses in Python (v3.9.12). For each pair of FIRE peaks, we calculated the expected co-accessibility as the product of their accessible proportions, while the observed co-accessibility was calculated as the proportion of fibers spanning both peaks that are accessible at each. We quantified the significance of this co-dependency using Fisher’s exact test. H3K27ac ChIP-seq (ENCFF365WHC, ENCFF522ANT, and ENCFF962HWM) and retinal DNase-seq (ENCFF064DIM) data was downloaded from ENCODE.

### Quantification of X chromosome and chromosome 13 allelic imbalance

To quantify chromatin allelic imbalance on each chromosome, we measured the number of accessible molecules (FIRE precision ≤ 0.10) for each haplotype at every FIRE peak on chromosomes 13 and X compared to the whole genome. We then calculated the difference in percent accessibility between the maternal and paternal haplotypes at every FIRE peak and built a distribution of these values to identify regions of imbalance. We tested for statistical significance by comparing the distributions of chromosomes X and 13 against the rest of the genome using a Mann–Whitney U test. FIRE peaks with unusually low coverage (coverage less than the median minus three standard deviations, or coverage less than ten reads) were excluded from the calculation to remove mapping, assembly, and phasing artifacts. Code to repeat this analysis is made available on GitHub as part of the Fiber-seq FIRE pipeline (https://github.com/fiberseq/fiberseq-fire, commit d095e67).

### Western blotting

Retinal organoid tissue was derived as described above from UDN318336 iPSCs (Day 72) and from AICS-0088 cl.83 iPSCs (Control sample, Day 66). Fibroblasts cell lines used were UDN318336, UND374570 (Control 1), and UND359892 (Control 2). Cells were lysed with RIPA lysis buffer supplemented with protease inhibitor tablets and processed as previously described^42^. Samples were normalized to 0.4-1 μg/μl, and 10 μl of sample was loaded on an SDS-PAGE gel. Antibodies (PDHA1: Abclonal A13687, p-PDHA1 S293: Abclonal AP1022, PDK3: Abclonal A8028, PDK3: Abclonal A12480, Beta Actin: Cell Signaling Technology 3700S) were used at 1:1000 dilution for Western blotting in 1% Bovine Serum Albumin (BSA) prepared in Tris-Buffered Saline with 0.1% Tween (TBS-T). HRP-conjugated secondary antibodies were used at 1:10.000 dilution and blots were imaged using iBrightCL1000. Quantification of western blotting was done using Fiji. The smallest rectangle possible was drawn around a band for a protein, and then was “measured” using FIJI’s measurement tool. This same sized rectangle was used to measure all other bands.

### Calcium uptake assay

Cells used were UDN318336 iPSCs and from AICS-0088 cl.83 iPSCs (Control). Calcium uptake assays were done as described before^42^. Cell number was determined using a Coulter counter. One million cells were used for each calcium uptake assay. Rate of calcium uptake was determined using data from the linear range of the uptake curves (40-60 sec.). Data from 3 replicates were normalized to the average of 3 control samples. A two tailed T-test was used for statistical analysis.

### Patient Consents

This study was approved by the National Institutes of Health Institutional Review Board (IRB) (IRB # 15HG0130), and written informed consent was obtained from all participants in the study.

## Acknowledgements

We would like to thank Christine C. Lambert, Aurelie Souppe, Harsharan Dhillon and Siyuan Zhang for assistance with library and sequencing preparations. A.B.S. holds a Career Award for Medical Scientists from the Burroughs Wellcome Fund and is a Pew Biomedical Scholar. This study was supported by National Institutes of Health (NIH) grants 1DP5OD029630 and 1U01HG010233, in addition to funds from the Collagen Diagnostic Laboratory, University of Washington, as well as funds from the Brotman Baty Institute for Precision Medicine. Sequence data analysis was supported by the University of Washington Center for Mendelian Genomics (UW-CMG), which was funded by NHGRI grant UM1 HG006493. The content is solely the responsibility of the authors and does not necessarily represent the official views of the National Institutes of Health. M.R.V. was supported by a training grant (T32) from the NIH (2T32GM007454-46).

## Data and materials availability

All GM12878 and GM23485 data is freely available through a public AWS bucket with direct links listed below. Restrictions apply to the availability of some data generated or analyzed during this study to preserve subject confidentiality. The corresponding author will on request detail the restrictions and any conditions under which access to some data may be provided. Cell lines obtained from the NIGMS Human Genetic Cell Repository at the Coriell Institute for Medical Research include GM12878 and GM23485. The UDN318336 fibroblast line was obtained directly from UDN participant UDN318336. Hia5 enzyme and plasmid is available upon request.

Fibertools BAM files for GM12878 and GM23485 aligned against GRCh38 are publicly available at the following aws links and can be used with IGV:

s3://stergachis-manuscript-data/2023/Vollger_et_al_long-read_multi-ome/GM12878_WGS.haplotagged.bam

s3://stergachis-manuscript-data/2023/Vollger_et_al_long-read_multi-ome/HG002_WGS.haplotagged.bam

Iso-seq BAM files for GM12878 and GM23485 aligned against GRCh38 are publicly available at the following aws links and can be used with IGV:

s3://stergachis-manuscript-data/2023/Vollger_et_al_long-read_multi-ome/isoseq/analysis-NA12878_bulkMAS-Seq-204367-mapped.bam

s3://stergachis-manuscript-data/2023/Vollger_et_al_long-read_multi-ome/isoseq/analysis-HG002_bulkMAS-Seq-204368-mapped.bam

Processed 5mC data in bigWig format for GM12878, GM23485, UDN318336, and retinal organoid derived from UDN318336 are publicly available at:

https://s3-us-west-1.amazonaws.com/stergachis-manuscript-data/index.html?prefix=2023/Vollger_et_al_long-read_multi-ome/5mC/

Processed FIRE results for GM12878, GM23485, UDN318336, and retinal organoid derived from UDN318336 are publicly available as trackhubs hosted on aws and can be directly linked into the UCSC genome browser:

https://s3-us-west-2.amazonaws.com/stergachis-manuscript-data/2023/Vollger_et_al_long-read_multi-ome/GM12878_pacbiome/trackHub/hub.txt

https://s3-us-west-2.amazonaws.com/stergachis-manuscript-data/2023/Vollger_et_al_long-read_multi-ome/HG002_pacbiome/trackHub/hub.txt

https://s3-us-west-2.amazonaws.com/stergachis-manuscript-data/2023/Vollger_et_al_long-read_multi-ome/UDN318336/trackHub/hub.txt

https://s3-us-west-2.amazonaws.com/stergachis-manuscript-data/2023/Vollger_et_al_long-read_multi-ome/UDN318336_retinal/trackHub/hub.txt

## Author Contributions

A.B.S, M.R.V., and J.K. conceived the overall method design, coordinated activities from co-authors, and wrote the first draft of the manuscript. M.R.V., E.S., and A.B.S., contributed to the analysis of Fiber-seq and MAS-Seq data. K.C.E., and T.A.R. contributed to the generation of patient-derived retinal organoids, as well as the evaluation of RNA-seq data in retinal organoids. Y.S. contributed to the western blots, as well as mitochondrial functional studies. Y.M., created the Hia5 enzyme. E.E.B., S.Sh., S.St., S.M.S., E.A.R., and A.M.W. contributed to the analysis of genomic data. U.S. and P.H.B. contributed to the Sanger validation of the translocation breakpoints. K.M.D., G.P.J., F.M.H.,M.J.B., A.L.Y., J.O., K.A.L., I.G., B.L.D., S.C., and A.A. contributed to patient phenotyping. M.H-P. contributed to getting IRB approval for this study. J.R., A.B.S, K.C.E., M.K., and Y.H.C. contributed to the generation of Fiber-seq and mRNA samples. J.G.U. contributed with the mRNA concatenation protocol development and transcriptome data analysis. C.T.S. and A.M.W. contributed to CpG methylation calling and IGV visualization. I.J.M, and K.M.M. coordinated and performed the sequencing runs. All authors contributed to the revision of the manuscript.

## Code availability

All software used in the study are publicly available.

## Conflicts

J.K., J.G.U., C.T.S., A.M.W., M.K. and I.J.M. are full-time employees at PacBio, a company developing single-molecule sequencing technologies. A.B.S. is a co-inventor on a patent relating to the Fiber-seq method (US17/995,058).

---

**Supplementary Table 1.**
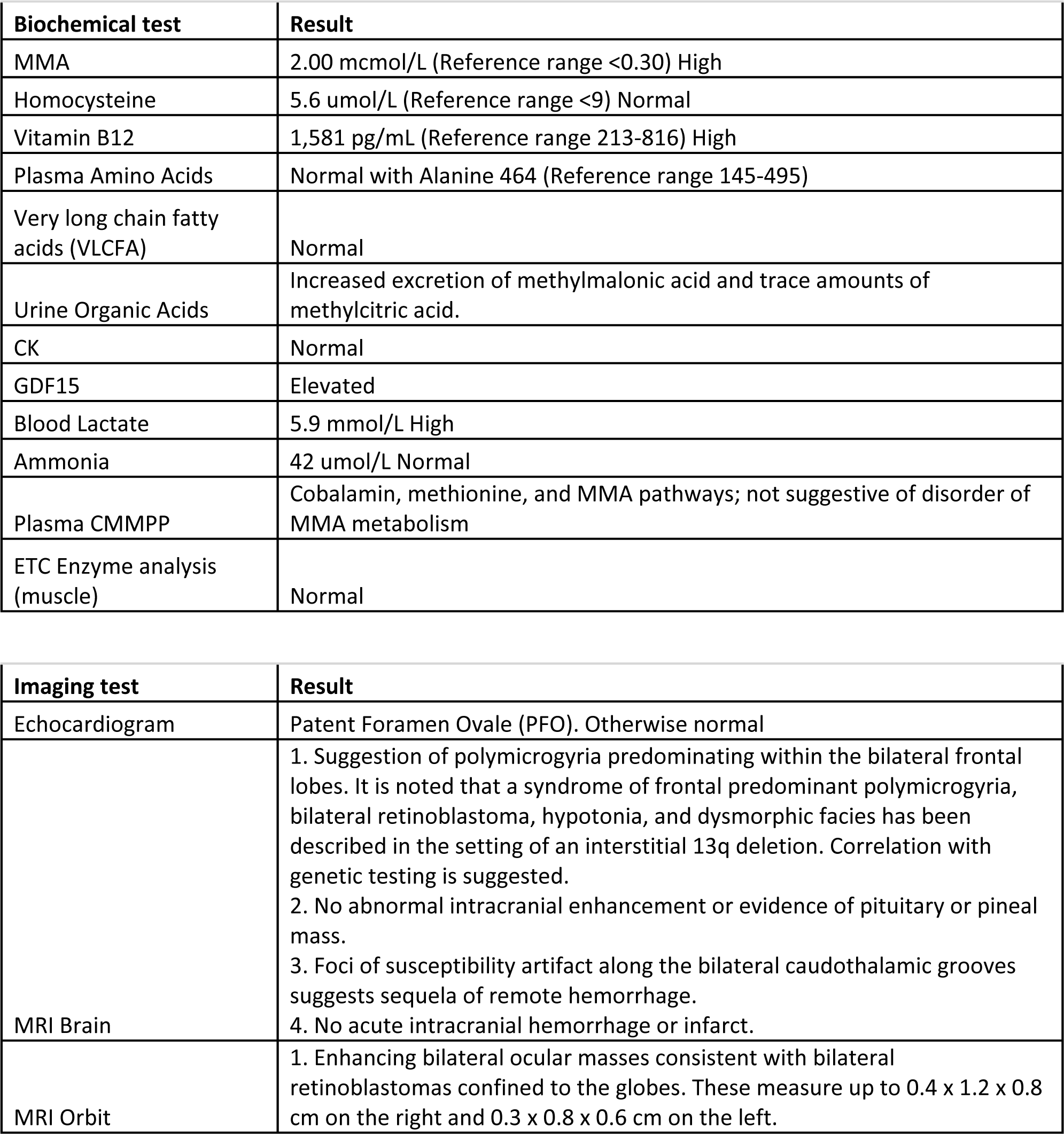
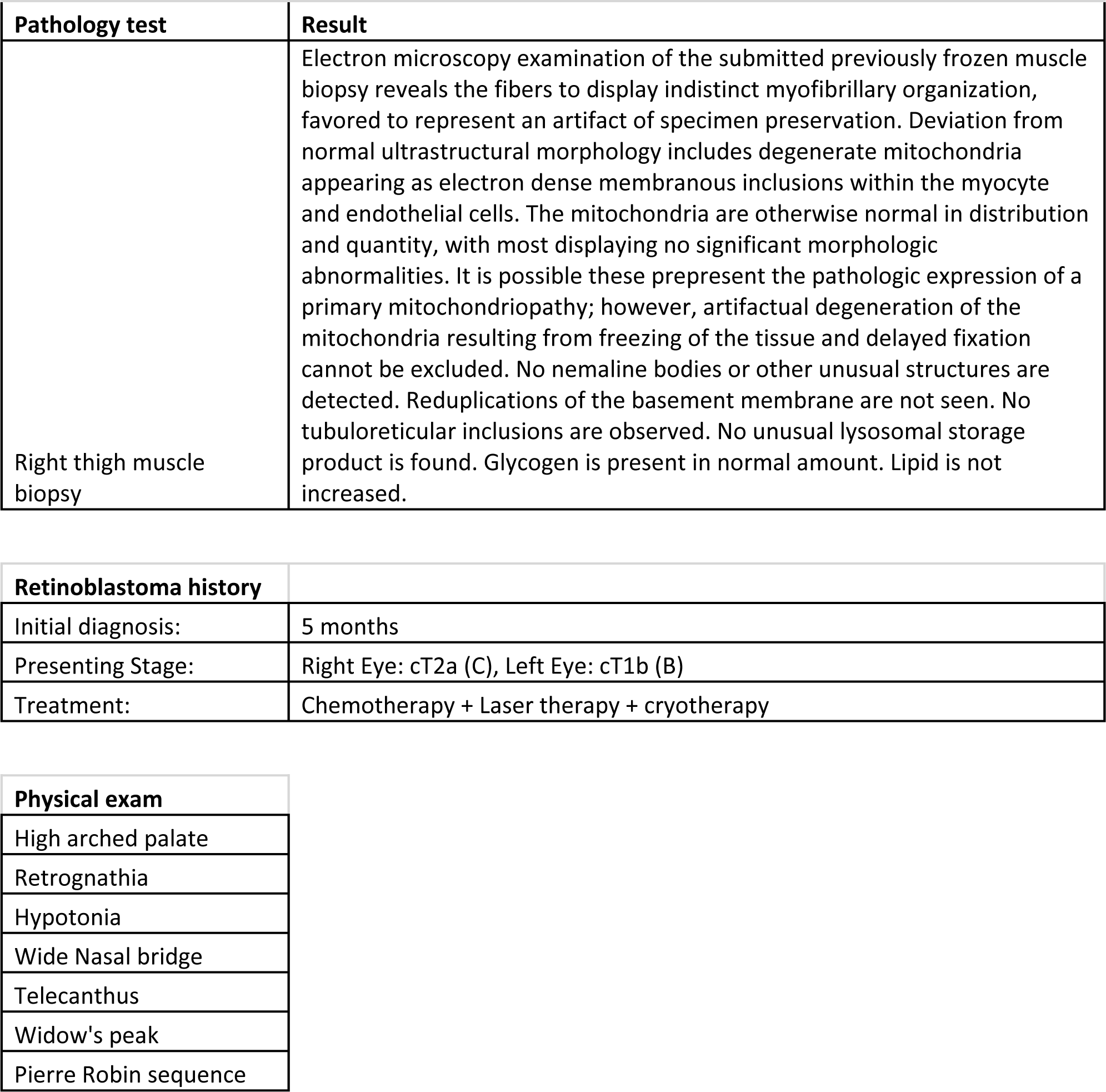
Clinical evaluation of proband.

**Supplementary Table 2.**
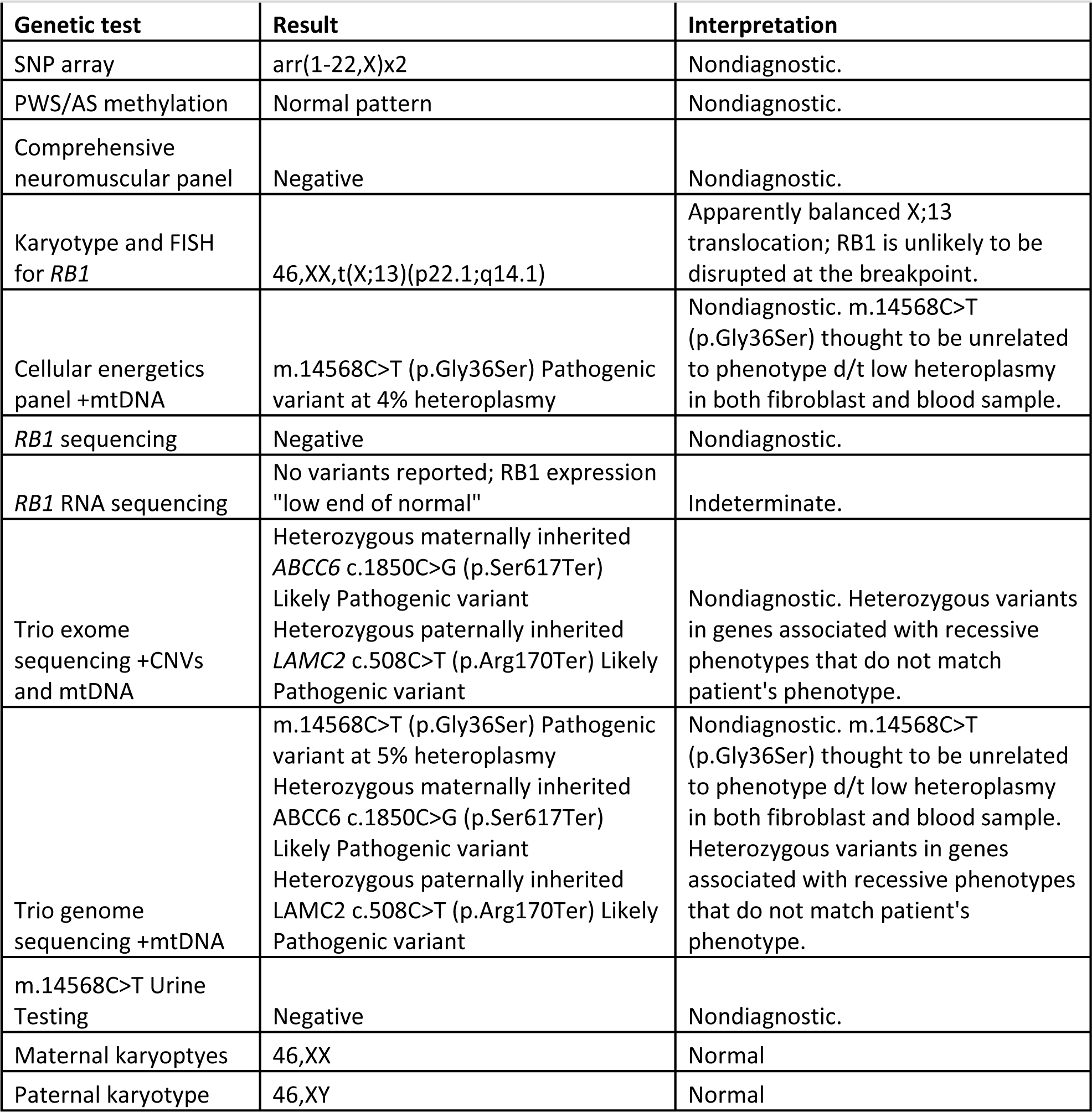
Summary of prior genetic evaluations for proband.

| Genetic test | Result | Interpretation |
| --- | --- | --- |
| SNP array | arr(1-22,X)x2 | Nondiagnostic. |
| PWS/AS methylation | Normal pattern | Nondiagnostic. |
| Comprehensive neuromuscular panel | Negative | Nondiagnostic. |
| Karyotype and FISH for *RB1* | 46,XX,t(X;13)(p22.1;q14.1) | Apparently balanced X;13 translocation; *RB1* is unlikely to be disrupted at the breakpoint. |
| Cellular energetics panel +mtDNA | m.14568C>T (p.Gly36Ser) Pathogenic variant at 4% heteroplasmy | Nondiagnostic. m.14568C>T (p.Gly36Ser) thought to be unrelated to phenotype d/t low heteroplasmy in both fibroblast and blood sample. |
| *RB1* sequencing | Negative | Nondiagnostic. |
| *RB1* RNA sequencing | No variants reported; *RB1* expression "low end of normal" | Indeterminate. |
| Trio exome sequencing +CNVs and mtDNA | Heterozygous maternally inherited *ABCC6* c.1850C>G (p.Ser617Ter) Likely Pathogenic variant<br>Heterozygous paternally inherited *LAMC2* c.508C>T (p.Arg170Ter) Likely Pathogenic variant | Nondiagnostic. Heterozygous variants in genes associated with recessive phenotypes that do not match patient's phenotype. |
| Trio genome sequencing +mtDNA | m.14568C>T (p.Gly36Ser) Pathogenic variant at 5% heteroplasmy<br>Heterozygous maternally inherited *ABCC6* c.1850C>G (p.Ser617Ter) Likely Pathogenic variant<br>Heterozygous paternally inherited *LAMC2* c.508C>T (p.Arg170Ter) Likely Pathogenic variant | Nondiagnostic. m.14568C>T (p.Gly36Ser) thought to be unrelated to phenotype d/t low heteroplasmy in both fibroblast and blood sample. Heterozygous variants in genes associated with recessive phenotypes that do not match patient's phenotype. |
| m.14568C>T Urine Testing | Negative | Nondiagnostic. |
| Maternal karyotypes | 46,XX | Normal |
| Paternal karyotype | 46,XY | Normal |

